# A non-canonical mechanism of GPCR activation

**DOI:** 10.1101/2023.08.14.553154

**Authors:** Alexander S. Powers, Aasma Khan, Joseph M. Paggi, Naomi R. Latorraca, Sarah Souza, Jerry Di Salvo, Jun Lu, Stephen M. Soisson, Jennifer M. Johnston, Adam B. Weinglass, Ron O. Dror

## Abstract

The goal of designing safer, more effective drugs has led to tremendous interest in molecular mechanisms through which ligands can precisely manipulate signaling of G-protein-coupled receptors (GPCRs), the largest class of drug targets. Decades of research have led to the widely accepted view that all agonists—ligands that trigger GPCR activation—function by causing rearrangement of the GPCR’s transmembrane helices, opening an intracellular pocket for binding of transducer proteins. Here we demonstrate that certain agonists instead trigger activation of free fatty acid receptor 1 by directly rearranging an intracellular loop that interacts with transducers. We validate the predictions of our atomic-level simulations by targeted mutagenesis; specific mutations which disrupt interactions with the intracellular loop convert these agonists into inverse agonists. Further analysis suggests that allosteric ligands could regulate signaling of many other GPCRs via a similar mechanism, offering rich possibilities for precise control of pharmaceutically important targets.

## Introduction

One-third of existing drugs act by binding to G protein–coupled receptors (GPCRs), and these receptors also represent the largest class of targets for the development of new therapeutics^1,2^. A tremendous amount of work has focused on understanding the molecular mechanism of GPCR activation—that is, how drugs and naturally occurring ligands cause GPCRs to adopt molecular conformations that stimulate intracellular signaling.

For decades, the dominant model of GPCR activation has been that agonists facilitate rearrangement of a GPCR’s seven conserved transmembrane (TM) helices^1,3–7^. These TM helices connect the extracellular and intracellular surfaces of the GPCR, allowing extracellular ligands to cause opening of a large intracellular pocket in which G proteins and other intracellular signaling proteins bind. Molecular structures, spectroscopic experiments, mutagenesis studies, and computer simulations involving many GPCRs and ligands have all supported this model, leading to the common assumption that all GPCR agonists act by facilitating rearrangement of the TM helices^8– 11^.

Most known GPCR ligands bind at the orthosteric site where endogenous ligands bind, but a recent explosion of GPCR structures has demonstrated that various ligands can bind to diverse sites across the GPCR surface^2,12–15^. Ligands that bind in such allosteric sites are of great interest for drug discovery, not least because they provide a mechanism to achieve high selectivity for target receptors^12,16^. Many allosteric GPCR ligands also stimulate or prevent receptor activation^14,17–21^. Like orthosteric ligands, allosteric ligands—including those that bind near the intracellular surface—are widely assumed to act by causing or preventing rearrangement of a GPCR’s transmembrane helices. Studies of multiple allosteric ligands have supported this model^18–20^.

We set out to investigate the detailed molecular mechanism by which AP8, an allosteric ligand that binds near the intracellular side of free fatty acid receptor 1 (FFAR1 or GPR40)^14,15^, activates this receptor (Fig. 1a). AP8 is a full agonist—it strongly stimulates activation of FFAR1 even in the absence of an orthosteric ligand. To our surprise, we discovered that AP8 acts via a mechanism fundamentally different from that of previously characterized allosteric and orthosteric GPCR ligands. Instead of favoring rearrangement of the transmembrane helices, AP8 changes the orientation of an intracellular loop, leading the receptor to couple more effectively with G proteins. Our results suggest that several other FFAR1 agonists also act via this mechanism, and that ligands could control signaling of many other GPCRs in a similar fashion.

**Figure 1.**
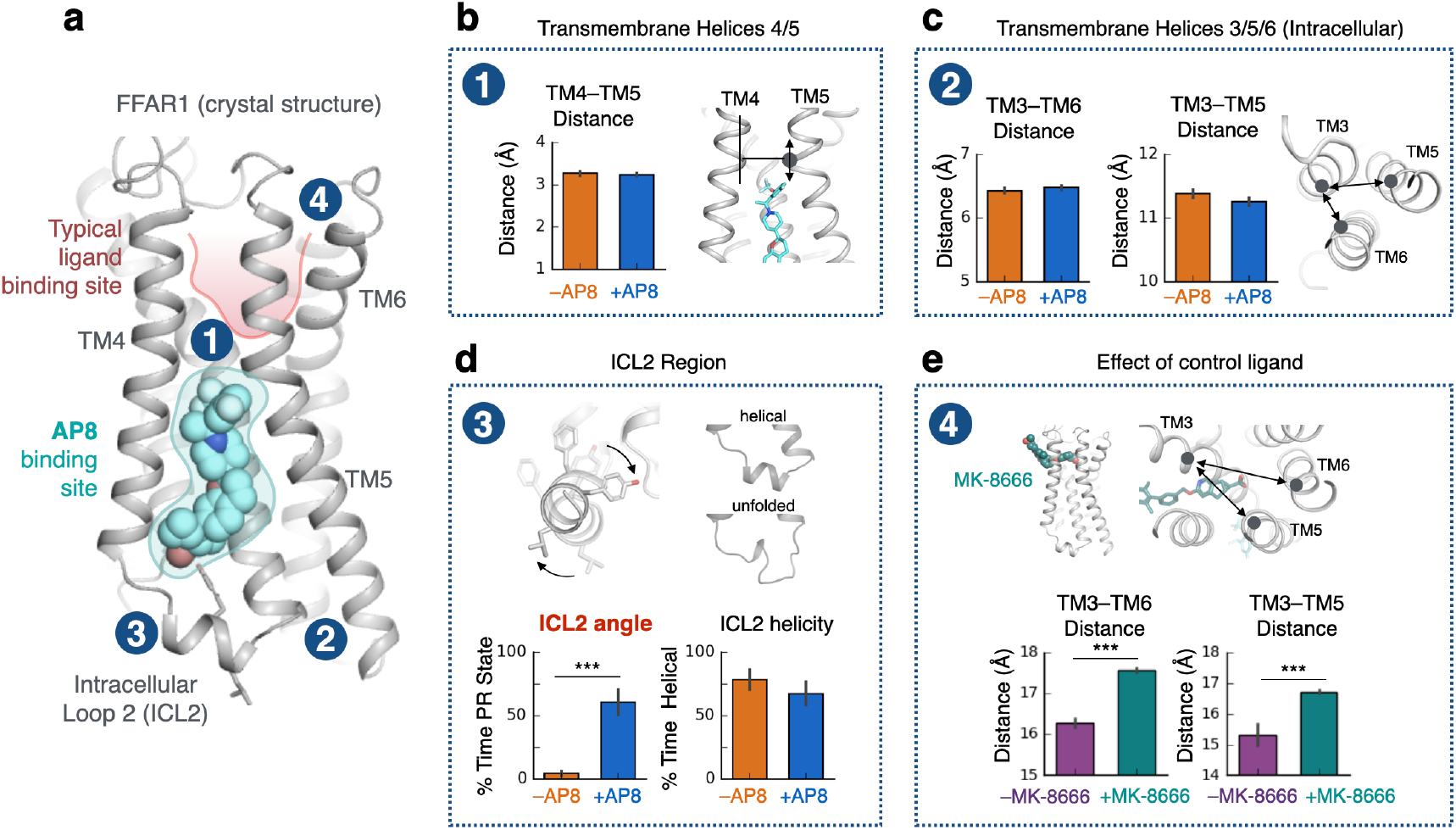
Unlike typical GPCR ligands, AP8 controls the conformation of intracellular loop but does not affect key transmembrane helices. **(a)** The crystal structure of FFAR1 (PDB 5TZY) shows agonist AP8 (cyan) bound to a membrane-facing pocket near intracellular loop 2 (ICL2) and transmembrane helix 3, 4, 5 (TM3, TM4, TM5). Other FFAR1 agonists, such as MK-8666, bind in a more typical pocket (highlighted in red) within the extracellular region of the receptor. **(b, c)** AP8 does not significantly affect the conformation of key TM helices in molecular dynamics simulations. The conformation of TM5 was measured by its vertical shift relative to TM4 (see Methods), which differs by 3 Å between the AP8-bound and AP8-free crystal structures. The intracellular conformation of TM helices was measured by distances between C? atoms (TM3-TM6: residue 105 C? to 222 C?, TM3-TM5: 104 C? to 208 C?). Data presented as mean of 5-10 independent simulations, each at least 2 µs in length; blue bars are simulations started from AP8-bound crystal structure and orange bars are simulations started from AP8-free crystal structure (n.s., not significant; P>0.05 for all comparisons by two-sided t-test; error bars are 68% confidence interval, CI). **(d)** AP8 has a significant effect on the orientation of the ICL2 helix in simulation as measured by the rotation about the helical axis (see Methods). The conformation of ICL2, both angle and helicity, was quantified over simulations with and without AP8 bound, started from the AP8-bound crystal structure. Data presented as mean of 5-10 independent simulations (*** indicates P<0.001 by two-sided t-test; error bars are 68% CI). **(e)** For comparison, control ligand MK-8666 does have a substantial effect on key TM helices, in particular the extracellular ends of TM3, 5, and 6 as measured by distances between C? atoms (TM3-TM6: residue 83 C? to 184 C?, TM3-TM5: 83 C? to 244 C?). Data presented as mean of 5-10 independent simulations; green bars are simulations started from MK-8666-bound crystal structure and purple bars are simulations with MK-8666 removed.

These findings indicate that the classical model of GPCR activation is incomplete: ligands can trigger activation not only by causing opening of the inter-helical transducer-binding pocket but also by directly rearranging intracellular receptor loops. Our results suggest a variety of opportunities for designing drugs that precisely target various GPCRs and provide fine control over their signaling.

### Allosteric agonist does not control transmembrane helix conformation

Previous hypotheses for the mechanism by which AP8 activates FFAR1 have been based on comparison of two crystal structures: one of FFAR1 bound to both AP8 and the orthosteric partial agonist MK-8666 (*AP8-bound structure*), and one of FFAR1 bound only to MK-8666 (*AP8-free structure*). These structures differ in two ways (Supp. Fig. 1, 2)^14^. First, TM5 is shifted 3 Å toward the extracellular end of the receptor relative to TM4 in the AP8-bound structure, along with small shifts in the intracellular ends of TM3 and TM6. Second, intracellular loop 2 (ICL2) is helical in the AP8-bound structure but unresolved in the AP8-free structure. These differences led to two hypotheses for the mechanism of agonism for AP8 and related ligands^14^: (1) these ligands may stabilize key transmembrane helices including TM5 and TM6 in the canonically “active” conformation that enables G protein binding, or (2) these ligands promote a helical ICL2 over a disordered ICL2 to directly stabilize part of the interface for G protein binding.

**Figure 2.**
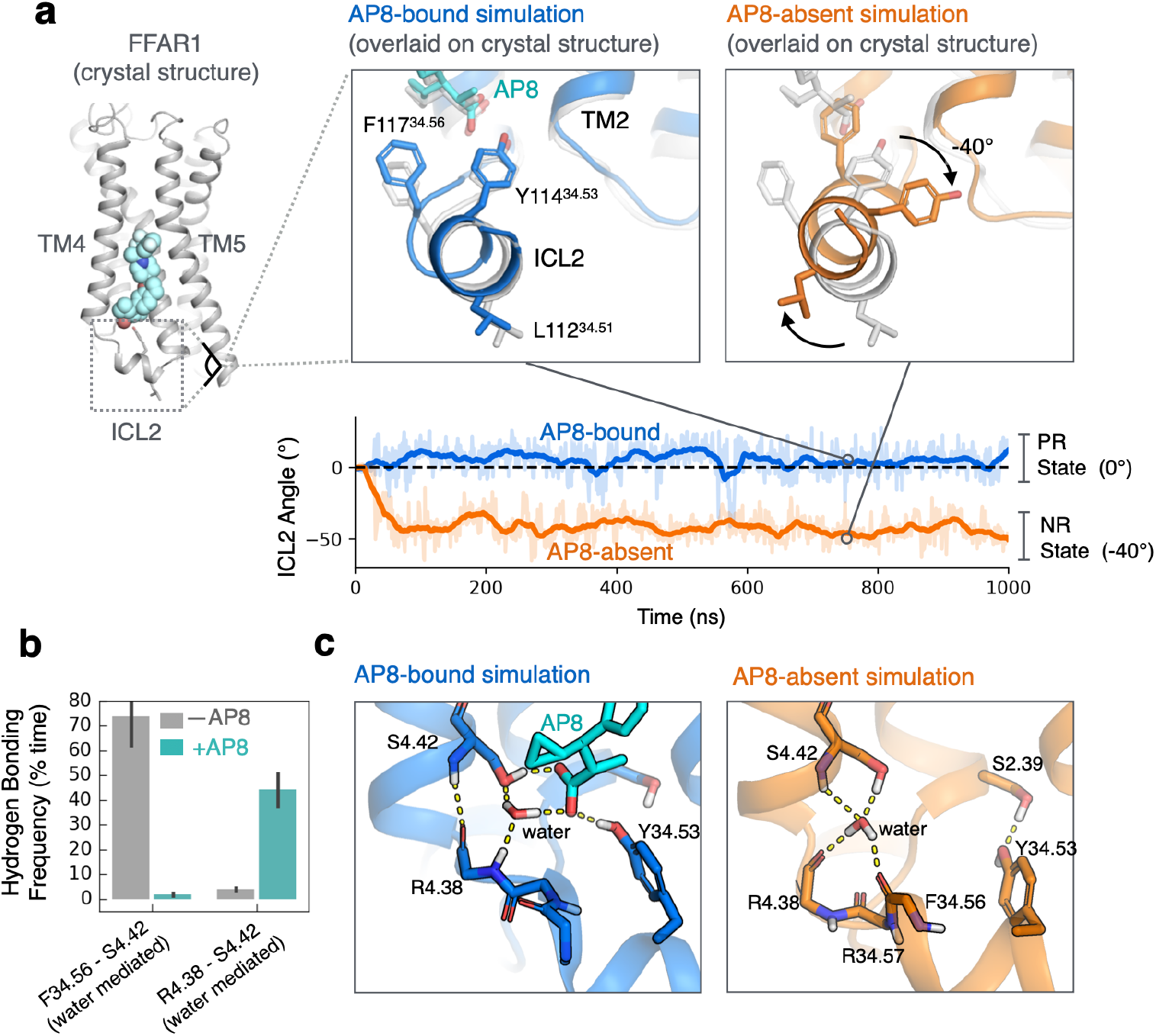
Ligand directly affects the intracellular receptor surface by rotating intracellular loop 2. **(a)** Simulations of the receptor with AP8 bound (blue) favor one stable ICL2 conformation (PR state) while simulations without AP8 (orange) favor a distinct negatively rotated ICL2 conformation (NR state). Both simulations were started from the same structure. Representative simulation snapshots are shown at indicated timepoints overlaid on the AP8-bound crystal structure (light grey). Simulation trajectories show the ICL2 angle as measured by the rotation of L112 and Y114 around the ICL2 helix axis (see Methods). Dashed line indicates the distance in the starting structure. **(b)** AP8 alters a network of polar interactions; the frequency of water-mediated hydrogen bonds between key ICL2 residues were quantified in presence and absence of AP8. Data presented as mean of 5-10 independent simulations (error bars are 68% CI). Only simulation frames where ICL2 remained folded were used. **(c)** Simulation frames show representative hydrogen bonds (yellow dashes) formed with and without AP8 bound. With AP8 bound, a stable water molecule forms a hydrogen bond network bridging AP8 and the ICL2 backbone. In the absence of AP8, the water molecule reorients to form a new stable network of hydrogen bonds, necessitating a rotation of ICL2.

To investigate these hypotheses, we performed extensive molecular dynamics (MD) simulations of FFAR1 in a hydrated lipid bilayer, with and without AP8 bound. We initiated simulations from the AP8-bound structure; we retained AP8 under one condition and removed it in the other. We also initiated simulations from AP8-free structure.

Strikingly, we observed that AP8 had little influence on the arrangement of transmembrane helices in these simulations (Fig. 1b, 1c, Supp. Fig. 1). In 2 µs simulations initiated from the AP8-bound structure, the removal of AP8 had minimal effect on the distances between TM helices. Moreover, in 2 µs simulations initiated from the AP8-free structure, the TM helices quickly, within 200 ns, adopted positions seen in the AP8-bound structure (Supp. Fig 2c). In other words, the differences between the AP8-bound and AP8-free crystal structures, including the 3 Å difference in TM5 position, did not persist in simulations and did not depend on the ligand. We found that the difference in TM3, TM5 and TM6 position between the crystals may be explained by a difference in crystal packing contacts around these helices (Supp. Fig. 2b).

By contrast, in simulations of FFAR1 with and without the orthosteric agonist MK-8666 bound, we observed substantial differences in the arrangement of transmembrane helices (Fig 1e, Supp. Fig. 1b). These motions are typical of those caused by GPCR agonists and provide a reference point for the atypical behavior of AP8^3,4^. We also note that in simulations with MK-8666 bound, AP8 has a stabilizing effect on the extracellular end of TM4 that contacts both ligands (Supp. Fig. 1a). This motion may underlie the observed binding cooperativity between the two ligands, but this is not a likely explanation for activation as AP8’s agonism is not dependent on MK-8666.

Because the AP8-bound and AP8-free crystal structures of FFAR1 represent inactive receptor states (no G protein bound), we further examined interactions between AP8 and the transmembrane helices in the active receptor state, with a G protein bound. In particular, we modeled and simulated the FFAR1-Gq-AP8 complex (see Methods). We then estimated interaction energies and calculated distances between AP8 and surrounding TM helix residues (Supp. Fig. 3b) in both active and inactive receptor states. There were no large differences in estimated interaction energies or distances between the active and inactive states, again suggesting that AP8 does not act by stabilizing the active-state conformation of the TM helices. Further supporting this conclusion, a recently published cryo-EM structure of the active-state FFAR1-Gq complex^22^ suggests that the TM helix residues immediately surrounding the AP8 binding site undergo very little conformational change upon activation (Supp. Fig. 3a).

### Allosteric agonist controls intracellular helix orientation

Next, we examined the alternative hypothesis that AP8 acts as an agonist by stabilizing ICL2 in a helical conformation. Our simulations did not support this hypothesis. First, in simulations initiated from the AP8-bound receptor structure, the ICL2 helix actually unfolds *faster* with AP8 bound than without (Fig 1d, Supp. Fig. 4). Second, using adaptively biased MD, we calculated the free energy difference between helical and disordered ICL2 conformations; the relative stability of the helical conformation did not differ significantly with and without AP8 bound (Supp. Fig. 5). Moreover, in several recent experimentally determined structures of GPCR-G_q_ complexes, ICL2 does not adopt a helical conformation, suggesting that a helical ICL2 conformation is not a requirement for G_q_ coupling^23,24^.

However, AP8 does cause one notable and robust conformational change in FFAR1 in simulation: AP8 controls the equilibrium between two distinct helical ICL2 conformations (Fig. 1d, 2a). In simulations initiated from the AP8-bound structure but with AP8 removed, the ICL2 helix rotates –40 degrees about its helical axis to a new conformation within nanoseconds (Fig. 2a, Supp. Fig. 4). This rotation hardly ever occurs with AP8 bound. We refer to the helical conformation in the AP8-bound structure as the positively rotated (PR) state, and to the helical conformation adopted upon removal of AP8 as the negatively rotated (NR) state. In additional simulations initiated with ICL2 in the NR state, the ICL2 helix rotated back to the PR state upon additional of AP8 (Supp. Fig. 6). These experiments support that AP8 controls the equilibrium between two well-defined ICL2 conformations. Additional control simulations with MK-8666 removed and with engineered mutations reversed showed consistent results (Supp. Fig. 7a).

How does AP8 trigger this rotation of the ICL2 helix? Simulations reveal that AP8 stabilizes a network of key polar interactions (Fig. 2b, 2c). First, a stable water molecule forms hydrogen bonds with both the AP8 carboxylate and the ICL2 backbone. This water is not modeled in the crystal structure, but there is consistent density at this location. In the absence of AP8, this water flips to form a different set of stable hydrogen bonds (Fig. 2c). Second, AP8 forms a frequent hydrogen bond with the sidechain of tyrosine 114(34.53) in the middle of the ICL2 helix. To test whether these hydrogen bonds explained the effect of AP8, we purposely disrupted these interactions in additional simulations by mutation or protonation (see Methods). These targeted disruptions prevented AP8’s effect, as ICL2 adopted the NR state even with AP8 bound (Supp. Fig. 7b). We thus concluded that AP8’s affect on the orientation of the ICL2 helix is due to the ligand’s ability to control this hydrogen bond network. We used this model to design the *in vitro* mutagenesis experiments discussed below.

### Intracellular helix orientation is coupled to G protein binding

Could a simple rotation of the ICL2 helix control G protein signaling? Structural analysis indicates that ICL2 fits neatly into a corresponding cavity on Gq when the ICL2 helix adopts the AP8-stabilized PR state, but not when the ICL2 helix adopts the NR state (Fig. 3a). Simulations of the full complex with and without AP8 bound further supported this analysis. We observed that binding of Gq alone—like binding of AP8—favors the PR ICL2 state (Fig. 3b). AP8 and Gq together stabilize this ICL2 conformation to an even greater degree. This implies that AP8 and Gq bind cooperatively to FFAR1: binding of AP8 increases the affinity of FFAR1 for Gq by causing ICL2 to adopt the PR state.

**Figure 3.**
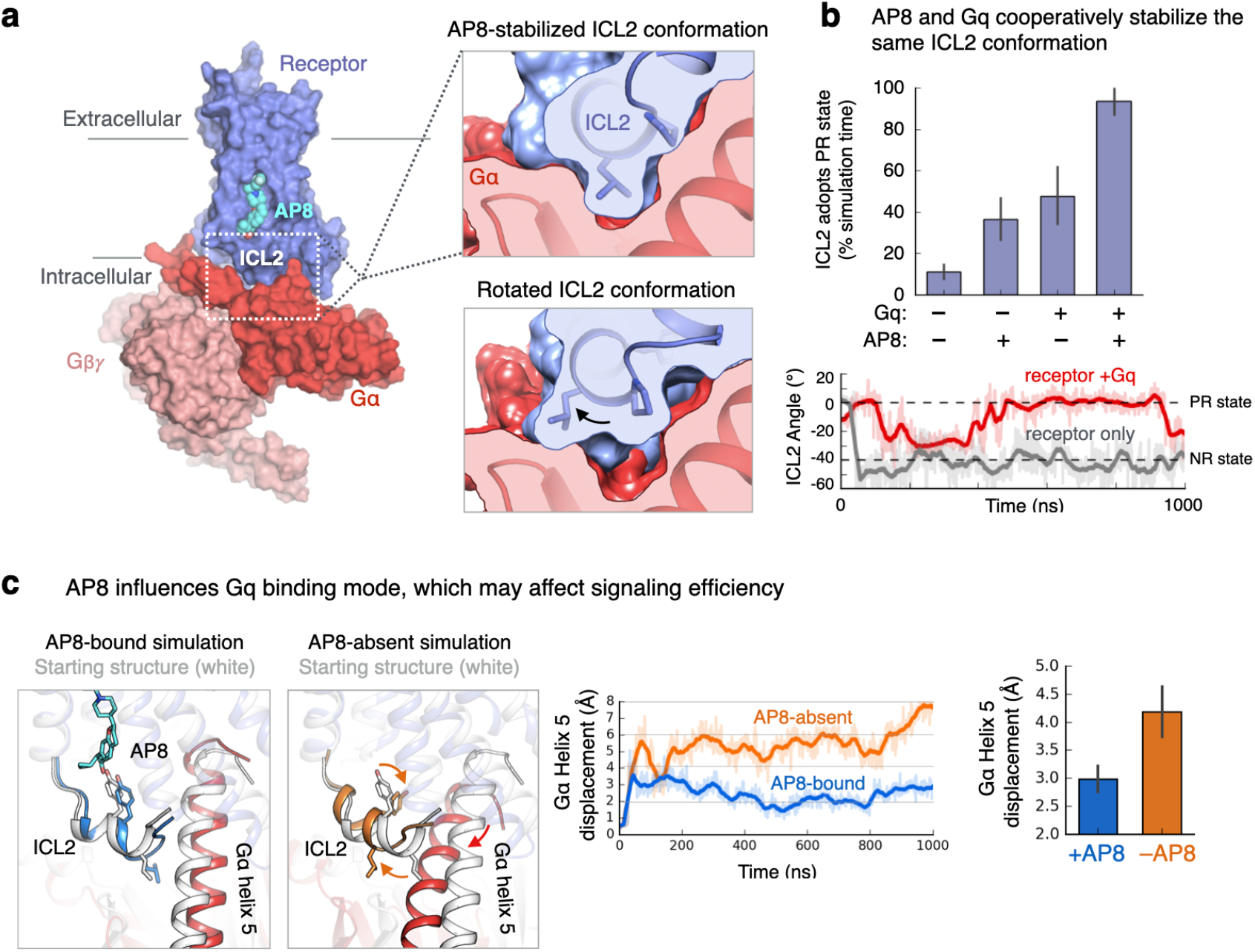
ICL2 helix orientation affects G protein binding. **(a)** Model of active FFAR1 with bound AP8 and heterotrimeric Gq constructed from homology modeling and alignment with other complexes (see Methods). Zoomed image shows that ICL2 in the AP8-stabilized conformation (PR state) forms a tight interface with Gq? (red). In the lower image, ICL2 modeled in the NR state has poor shape complementarity with the Gq surface. **(b)** AP8 and Gq both independently stabilize the PR state of ICL2, and do so to an even greater degree together, indicating a cooperative effect. Data presented as mean of 5-10 independent simulations for each condition, each 1 µs in length (error bars are 68% CI). The simulation trace below shows ICL2 angle vs. time for the receptor only condition (grey) and receptor-Gq condition (red). When Gq is bound to the receptor, the ICL2 PR state is stabilized relative to the receptor alone. **(c)** ICL2 conformation is also coupled to the orientation of Gq relative to the receptor, in particular Gα helix 5. Representative simulation frames (left) and traces (middle) of the FFAR1-Gq model are shown with and without AP8 bound. In images, the starting structure is shown in grey and the simulation frame in color. The displacement of the Gα helix 5 was calculated by aligning simulation frames on the receptor and calculating the root mean square displacement (RMSD) of helix 5 (terminal 10 residues) relative to the starting structure. Bars (right) show mean displacement from 5-10 independent simulations for each condition, each 1 µs in length (error bars are 68% CI).

In addition to modulating the binding affinity of FFAR1 for the G protein, rotation of the ICL2 helix might influence the signaling properties of the FFAR1-G protein complex. Previous studies have indicated that, in other GPCRs, certain ICL2 mutations prevent G protein activation (i.e., GDP-GTP nucleotide exchange) without preventing G protein binding to the receptor^25,26^. This suggests that the ICL2 interface is important for the exchange of GDP for GTP within the G protein. Indeed, when comparing simulations of the FFAR1-Gq complex with and without AP8 bound, we found that the presence of AP8 leads to a change in the orientation of the Gq α5 helix relative to both the FFAR1 and the rest of Gq (Fig. 3c, Supp. Fig. 9). Because the α5 helix extends from FFAR1 to the nucleotide binding site in Gq, such a change could potentiate nucleotide exchange.

### Experimental validation of non-canonical activation mechanism

To validate our molecular mechanism for ICL2-mediated signaling, we conducted several *in vitro* experiments. First, a key component of our proposed mechanism is that AP8’s carboxylate group forms polar interactions that hold ICL2 in a conformation favorable for G protein signaling (Fig. 2). We reasoned that a mutation to FFAR1 that introduces a carboxylate in a similar position could increase constitutive activity of the receptor, effectively mimicking AP8’s interactions. We chose to mutate glycine 3.49(103), which is directly adjacent to the AP8 carboxylate binding site, to glutamate. Simulations of the G3.49E mutant showed the carboxylate sidechain of E3.49 indeed often forms the same interactions as the AP8 carboxylate (Fig. 4c). IP1 accumulation assays in HEK293 cells expressing FFAR1 were used as a measure of Gq-mediated signaling. The G3.49E mutation significantly increased basal activity (normalized for receptor surface expression) in agreement with the proposed mechanism (Fig. 4d, Supp. Fig. 10c, Supp. Table 1)^27^. By contrast, mutating G3.49 to residues that could not form these hydrogen bonds—including aspartate, whose carboxylate does not extend far enough, as well as leucine and alanine—decreased basal activity. Our computationally derived mechanism elegantly explains this trend, which is otherwise surprising when compared to the behavior of other class A GPCRs. Most class A GPCRs have a carboxylate residue (aspartate or glutamate) at position 3.49, which stabilizes the inactive state; mutating it to a neutral residue typically increases basal activity^28,29^. Our finding that FFAR1’s basal activity instead increases when one replaces a neutral residue at this position with glutamate supports our proposed mechanism and underscores the significance of the unique polar network we observed at FFAR1.

**Figure 4.**
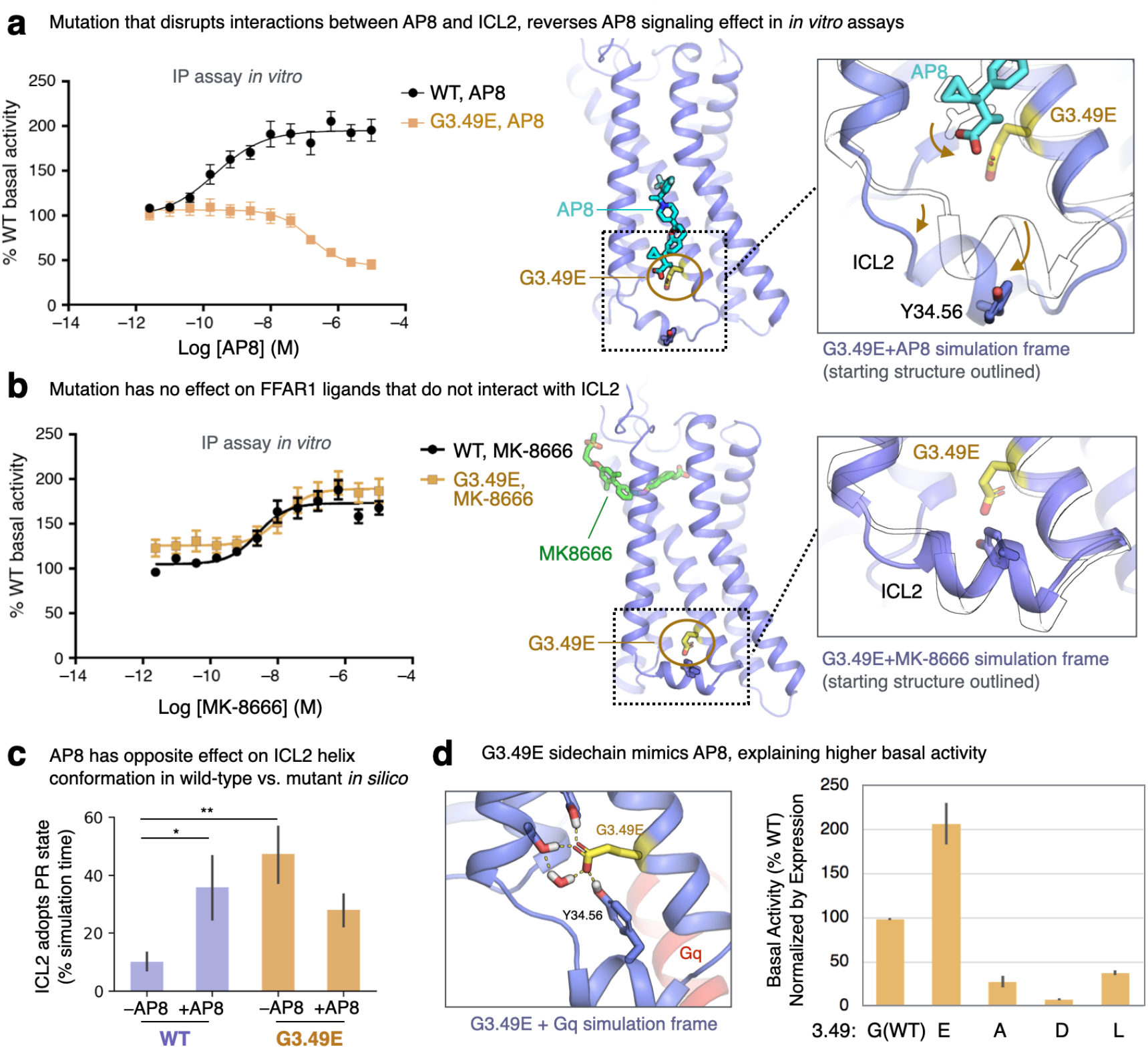
Mutagenesis experiments validate computationally derived activation mechanism. **(a, b)** At the mutated G3.49E receptor, AP8 acts as an inverse agonist. FFAR1 activity was monitored in IP1 accumulation assays in HEK293 cells expressing WT or G3.49E mutant receptors treated with (a) AP8 or MK-8666. Data is plotted as the % of WT receptor basal activity (cells treated with 1% DMSO), where data points are mean ± S.E.M. from N=3 experiments. Dose response curves were fit to a standard 4 parameter non-linear regression model. Images at right show simulation frames in color and starting structure as a black outline; AP8 is displaced from its WT binding pose by G3.49E, likely due to repulsion of the two nearby carboxylates. **(c)** In simulations, AP8 destabilizes the PR ICL2 state at the G3.49E receptor, the opposite of its behavior at the WT receptor. Data presented as mean of 5-10 independent simulations for each condition, each 1 µs in length (error bars are 68% CI). **(d)** Basal receptor activity in the IP1 accumulation assay, normalized by receptor surface expression, is plotted at right for different mutants at the 3.49 position. Only G3.49E leads to an increase in activity relative to WT. At left, a snapshot of the simulation of the G3.49E receptor in complex with Gq shows the glutamate sidechain can mimic the interactions of the AP8 carboxylate.

Second, we observed that, whereas AP8 increases the fraction of time that ICL2 spends in the PR state in simulation of the wild-type FFAR1, AP8 has the opposite effect at the G3.49E mutant (Fig. 4c). In simulations of the mutant, AP8’s carboxylate tends to be pushed outward, away from the glutamate, disrupting the hydrogen bond network that otherwise stabilizes ICL2 in the PR state (Fig. 4a). Indeed, our in vitro experiments show that AP8 acts as an *inverse agonist* at the G3.49E receptor; AP8 lowered the IP1 accumulation levels to 40 ± 6% of basal activity. Likewise, the G3.49E mutation converts AP3—an analogue of AP8 that binds at the same membrane-facing site—from an agonist to an inverse agonist (Supp. Fig. 10a). In contrast, the G3.49E mutation does not alter the efficacy of orthosteric agonist MK-8666, which remains an agonist at the mutant receptor (Fig. 4b, Supp. Table 2).

Another key interaction between AP8 and ICL2 is the hydrogen bond to Y114(34.53). Simulations showed that ICL2 mutation Y114(34.53)F diminishes AP8’s ability to control ICL2, though it does not reverse its effect (Supp. Fig. 7d). As predicted, Y114F decreased the potency of AP8 by 700x in IP1 accumulation assays despite only causing a 15x decrease in affinity, indicating reduced efficacy (Supp. Fig. 10d, Supp. Table 1). The effect on MK-8666 was minimal in comparison (Supp. Table 2).

An unexpected aspect of our mechanism is that AP8-like ligands do *not* act by stabilizing a single globally active receptor state through rearrangements of transmembrane helices. Using the G3.49E mutant, we could further test this aspect of our model. If AP8-like ligands, like MK-8666, simply acted on the transmembrane helix bundle to stabilize either inactive or active receptor states, then an inverse agonist would be predicted to have negative binding cooperativity with an agonist. Thus, based on the ternary complex model, one would expect AP8-like ligands to behave as negative allosteric modulators (NAMs) at the G3.49E construct. Instead, as our model predicts, the positive binding cooperativity between AP8-like ligands and MK-8666 is maintained at the G3.49E mutant despite the fact that the ligands have opposing functional effects (Supp. Fig. 10b). This supports our mechanism; AP8-like ligands control receptor activation through ICL2 independent of other effects on the receptor that may contribute to ligand binding cooperativity. We note that in simulations with MK-8666 bound, AP8 has a stabilizing effect on the extracellular end of TM4 that contacts both ligands (Supp. Fig. 1a) and indeed the presence of AP8 is associated with reduced dynamics of MK-8666. This motion may underlie the observed binding cooperativity between the two ligands.

As an additional test of our conclusion that AP8 acts not by stabilizing a helical ICL2 over a disordered ICL2 but by stabilizing one helical ICL2 conformation over another, we made a helix-destabilizing mutation of alanine to glycine at position 34.55(116), a residue located in the ICL2 helix but not directly contacting AP8. This mutation slightly increased AP8’s E_max_ relative to wild type (Supp. Table 1). This is consistent with our simulation findings that stabilizing a folded over disordered ICL2 helix is not crucial to AP8’s agonism.

### ICL2 mechanism generalizes beyond one ligand or receptor

How generalizable is ICL2-mediated signaling? To address this question, we turned to other ligands and class A GPCRs. An analysis of available GPCR structures showed diverse molecules binding around ICL2 and within the groove formed by TM3/4/5, especially lipids and sterols (Fig. 5c). Fatty acids and other lipids, being membrane-soluble, are well positioned to interact with this membrane-facing pocket. Along these lines, it has been proposed that endogenous fatty acids interact with the ICL2 site in FFAR1, though possibly with low affinity^14,15,30^. To explore this possibility, we conducted 2 µs simulations of a long-chain fatty acid, linolenic acid (ALA), modeled into the AP8 binding site (Supp. Fig 11). ALA remained stably bound to the site in all simulations and favored the same ICL2 conformation as AP8 through similar interactions (Supp. Fig 11). Though a deeper study of receptor modulation by fatty acid is beyond the scope of this work, these results support the potential of diverse molecules to regulate this site.

**Fig 5.**
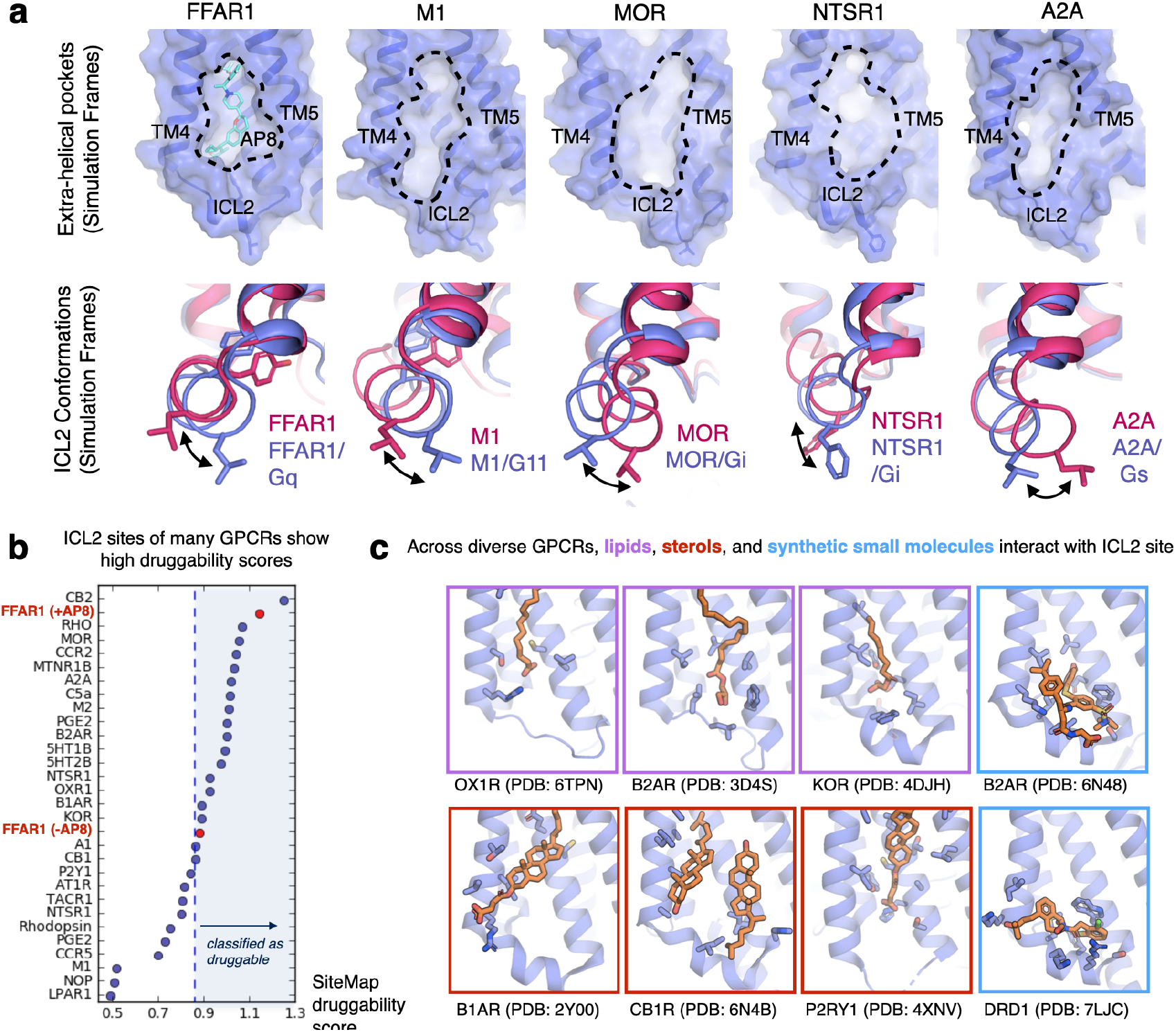
Druggable allosteric pockets exist at the same membrane-facing site in diverse GPCRs. **(a)** Selected frames from simulations of a range of different GPCRs show comparable allosteric pockets to the AP8 binding pocket. All receptors are shown from the same view angle with the putative pockets highlighted by dashed lines. Below, selected simulation frames show changes in the ICL2 helix angle and orientation upon formation of receptor-G protein complex, suggesting functional relevance. Purple structures were selected from simulations of receptor-G protein complexes, red structures were selected from simulation of receptor only. **(b)** Using Schrodinger’s SiteMap software, we scored the druggability of ICL2 allosteric sites from 29 class A GPCRs. Representative static structures from the Protein Data Bank were used. Druggable classification is determined from previously reported literature benchmarks. Curated structures from the PDB, with non-protein molecules bound to the ICL2 site. Receptors are shown in purple with ligands and other molecules in orange sticks.

Beyond FFAR1, only a handful of synthetic small molecules bind to the ICL2 site at present, but a structural analysis suggests broad opportunities. Compound 6 at B2AR and Mevidalen (LY3154207) at the D1 receptor bind directly above ICL2; these ligands act as both PAMs for orthosteric agonists and weak agonists on their own^21,31–33^. To more quantitatively assess the druggability of this site, we analyzed prospective ICL2 allosteric sites and their predicted propensity for ligand binding using 29 class A GPCR structures selected for diversity and a well-resolved ICL2 region. We used the FFAR1 structures with and without AP8 as references. Schrodinger’s SiteMap software was used to score druggability, which reflects the volume, curvature, and polarity of a potential binding site (Fig. 5b). Out of 29 GPCRs, the majority had ICL2 allosteric pockets with scores in the range empirically predicted to be druggable^34^. Several candidates, such as the mu-opioid (MOR) and cannabinoid 2 receptor (CB2R), had comparable scores to the AP8-bound FFAR1 structure.

These observations of putative pockets adjacent to ICL2 suggest that ligands could be designed to specifically bind to these sites, but would such ligand be able to modulate signaling? Using our existing library of GPCR simulations, we analyzed this site and dynamic ICL2 conformations for muscarinic (M2), mu-opioid (MOR), neurotensin (NTSR1), and adenosine (A2A) receptors. These are key targets for treatment of pain, addiction, Parkinson’s disease, Alzheimer’s, schizophrenia, and cancer^3,35–37^. We found that clear pockets within the cleft of TM4 and TM5, comparable to the AP8 pocket, form in these other GPCRs (Fig. 5a). Like at FFAR1, we found in simulations that ICL2 helix angle and orientation differs with and without G-proteins bound to the receptor (Fig. 5b). To show these changes quantitatively, we analyzed the principal components of the ICL2 dihedrals during these simulations (Supp. Fig. 12). Thus, just as in FFAR1, ligands at this site could control signaling by stabilizing particular ICL2 conformations and consequently affecting the receptor interaction with G-proteins or other signaling partners like arrestins.

## Discussion

In summary, our results demonstrate that ligands can stimulate receptor signaling by precisely controlling the conformation of an intracellular loop. This is fundamentally different from the canonical mechanism utilized by the vast majority of GPCR agonists, which primarily involves rearranging key transmembrane helices and switches within the receptor core. Instead, using both atomic simulations and *in vitro* experiments, we showed that intracellular loop conformation alone can have a significant effect on G-protein binding and signaling. This also differs from the proposed mechanisms of other recently discovered allosteric ligand that binds near the same site, such as compound 6 at B_2_AR. Compound 6 was hypothesized to function by stabilizing ICL2 as a helix which then *triggers* a rearrangement of TM3 and TM6 to stabilize the canonical active state of the receptor. Overall, our mechanism provides fundamental insight for the design of future drugs that target GPCR allosteric sites.

By studying other GPCRs, we found that this same binding site and mechanism could likely be utilized at many receptors with exciting implications for future therapeutics. Indeed, the recent discovery of other agonist-PAMs that bind near ICL2 at other receptors, suggests this activation mechanism is not limited to FFAR1^21,33^. The ICL2 site may offer opportunities for greater selectivity, pathway bias, and allosteric modulation of orthosteric ligands. However, these sites are shallow and membrane-facing; finding functional, high-affinity ligands that bind to such novel allosteric sites is an important challenge for future efforts. With enhanced knowledge of structure and mechanism, more targeted structure-based approaches may be possible. For example, virtual screening by docking large small-molecule libraries to this particular site may be able to find ligands that bind here more efficiently. By leveraging unique chemical properties of known ligands that bind near ICL2, we can focus the screening efforts on chemically similar compounds that have a higher chance of being functionally relevant. Using frames from MD simulation, such as those illustrated in Fig. 5, we could better select receptor conformations where allosteric pockets are maximally accessible and ICL2 is in a desired state (e.g. bound to a G protein or arrestin). This approach could extend to other putative intracellular allosteric sites as well, such as proximal to ICL1 or ICL3^38^.

Although we focused on G proteins in this study, ICL2 is a crucial interface for both G proteins and arrestins^39,40^. The design of biased drugs that cause GPCRs to favor or avoid stimulation of arrestins relative to G proteins could lead to more effective treatments for a wide range of diseases^41^. Several lines of evidence suggest the ICL2 allosteric site is a promising opportunity to develop such biased ligands. Previous work by our lab showed that the ICL2 conformation is particularly relevant for arrestin activation^40^. Moreover, ICL2 can adopt different conformations when binding to arrestins vs. G protein^42^. In functional studies of the 5HT2A receptor, mutation of ICL2 residues have been shown to dramatically alter arrestin binding relative to G protein binding, enhancing pathway bias^43^. Thus, ligands that stabilize different ICL2 conformations would potentially discriminate between these signaling pathways^44^. Future structural work will be needed to examine how ICL2 conformation differs when bound to arrestins and different families of G proteins across a wider variety receptors.

Our results point toward a new and widely applicable direction for GPCR drug design. By exploiting the conformational heterogeneity of ICL2 and its role in signaling, this site could offer rich new possibilities for control of GPCR function in many pharmaceutically relevant targets.

## Acknowledgments

Funding was provided by a National Science Foundation Graduate Research Fellowship (A.S.P.), the Stanford ChEM-H Chemistry/Biology Interface Predoctoral Training Program under NIH Award Number T32GM120007 (A.S.P.), and a Stanford Graduate Fellowship (J.M.P.)

## Author Contributions

A.S.P and J.M.P. setup and performed molecular dynamics simulations. A.K., S.M.S., J.D.S., and J.L. performed mutagenesis, ligand binding, and activity experiments. A.S.P., J.M.P, and N.R.L. analyzed simulations and developed structural analyses. R.O.D., J.M.J., and A.M.W. supervised the project. A.S.P., A.K., and R.O.D. wrote the paper with input from all authors.

## Material and Methods

### Running Simulations

29 simulation conditions with 185 individual simulations were investigated as listed in Supp. Table 4. For simulation conditions 1-16, the AP8-bound crystal structure of FFAR1 (PDB 5TZY) was used as the starting point. The structure for these simulations was prepared by first removing the co-crystallized T4 lysozyme. Prime (Schrodinger) was used to model in missing side-chains and missing extracellular and intracellular loops. The thermostabilizing point mutations were maintained unless specified. For conditions 10-16, 20, and 21 mutations were introduced. Sidechains were mutated using Maestro (Schrodinger) and Maestro’s rotamer library was used to select an initial rotamer that best minimized clashes. For conditions 6-7, ICL2 was first removed and remodeled using an ICL2 segment from condition 2, where ICL2 had adopted the NR state. For condition 17, the AP8-absent crystal structure of FFAR1 (PDB 5TZR) was used and prepared similarly to above. For conditions 18-21, we used homology modeling in Prime to build an active-state model of FFAR1 in complex with heterotrimeric Gq. For modeling the active FFAR1 receptor, we used a composite modeling approach. For TM1-4 and connecting loops, with the exception of the DRY motif on TM3, we used the FFAR1 structure 5TZY as a template. This allowed preservation of the AP8 binding site. TM5-7 and the DRY motif used B2AR (PDB 3SN6) as a template. The alignment of FFAR1 and B2AR sequences was automatically generated, and then corrected to ensure that all helical domains were accurately modeled as per the 5TZY structure. For modeling Gq, we used templates from a related GPCR G11 complex (PDB:6OIJ) and GPCR Go complex (PDB:6G79). All template structures were first aligned to TM1-4 of the receptor. Palmitoylations were added to the N-terminus of Gq^45^. Both the active receptor and Gq model were constructed simultaneously in a single Prime model to ensure a clean interface.

For all FFAR1 simulations, interior waters were added from the higher resolution FFAR1 crystal structure 4PHU. AP8’s carboxylate was left deprotonated (except in condition 5), in accordance with its low pKa, solvent accessibility, and necessity to form hydrogen bonds with surrounding h-bond acceptors. The prepared protein structure was aligned on the transmembrane helices to the Orientation of Proteins in Membranes (OPM) structure of PDB entry 4PHU^46^. Parameters for AP8, MK-8666, and ALA were generated using the CHARMM General Force Field (CGenFF) with the ParamChem server^47^. Full parameter sets are available upon request. Across all multi-microsecond simulations, the ligands remained stably bound within the binding pocket and formed persistent contacts with surrounding residues.

For conditions 22-29, the PDB structure described in Supp. Table 4 was downloaded and prepared using the relevant protocols described above.

For all simulations, hydrogen atoms were added, and protein chain termini were capped with neutral acetyl and methylamide groups. PropKa was used to determine the dominant protonation state of all titratable residues at pH 7^48^. The Dowser program was used to hydrate any additional pockets within and around the GPCR. Then the receptor was inserted into a pre-equilibrated palmitoyl-oleoyl-phosphatidylcholine (POPC) bilayer using Dabble^49,50^. Sodium and chloride ions were added to neutralize each system at a concentration of 150 mM. Approximate system dimensions were 80 Å x 90 Å x 85 Å for receptor-only simulations, and 120 Å x 120 Å x 140 Å for receptor G-protein complexes. We used the CHARMM36 parameter set for protein molecules, lipids, and ions, and the CHARMM TIP3P water model for waters^51,52^.

All simulations were run on a single Graphical Processing Unit (GPU) using the Amber18 Compute Unified Device Architecture (CUDA) version of particle-mesh Ewald molecular dynamics (PMEMD)^53^. For each independent simulation, the system was minimized with 500 steps of steepest descent followed by 500 steps of conjugate gradient descent three times. 10 and 5 kcal mol^-1^ Å^-2^ harmonic restraints were used on the protein, lipid, and ligand atoms for the first and second minimization, respectively. 1 kcal mol^-1^ Å^-2^ harmonic restraints were used on the protein and ligand atoms for the final minimization. The systems were then heated over 12.5 ps from 0 K to 100 K in the NVT ensemble using a Langevin thermostat with harmonic restraints of 10.0 kcal·mol^-1^·Å^-2^ on the non-hydrogen atoms of the lipids, protein, and ligand. Initial velocities were sampled from a Boltzmann distribution. The systems were then heated to 310 K over 125 ps in the NPT ensemble. Equilibration was performed at 310 K and 1 bar in the NPT ensemble, with harmonic restraints on the protein and ligand non-hydrogen atoms tapered off by 1.0 kcal·mol^-1^·Å^-2^ starting at 5.0 kcal·mol^-1^ ·Å^-2^ in a stepwise manner every 2 ns for 10 ns, and finally by 0.1 kcal·mol^-1^·Å^-2^ every 2 ns for an additional 18 ns. All restraints were completely removed during production simulation. For standard molecular dynamics (all conditions except 8 and 9), production simulations were performed at 310 K and 1 bar in the NPT ensemble using the Langevin thermostat and Monte Carlo barostat. The simulations were performed using a timestep of 4.0 fs while employing hydrogen mass repartitioning^54^. Bond lengths were constrained using SHAKE. Non-bonded interactions were cut off at 9.0 Å, and long-range electrostatic interactions were calculated using the particle-mesh Ewald (PME) method with an Ewald coefficient (β) of approximately 0.31 Å and B-spline interpolation of order 4. The PME grid size was chosen such that the width of a grid cell was approximately 1 Å. Snapshots from each trajectory were saved every 200 ps during the production phase of each simulation. The AmberTools18 CPPTRAJ package^55^ was used to reimage trajectories, while Visual Molecular Dynamics (VMD)^56^ and PyMol (Schrodinger) were used for visualization and analysis.

For simulation conditions 8 and 9, we employed adaptively biased molecular dynamics (ABMD) in Amber^53^. Specifically, we used flooding mode, with a flooding timescale of 200 picoseconds and monitor frequency of 5000 picoseconds. All other production settings were the same as those previously described. We defined a multi RMSD collective variable using the backbone nonhydrogen atoms of ICL2 residues 111-118. For sampling free energy along this collective variable, we specified a resolution of 0.2, minimum of 0, and maximum of 5.5. The free energy along this coordinate was then collected in a separate output file.

### ICL2 Conformational Analysis

ICL2 helicity was determined by measuring the fraction of backbone hydrogen bonds between the i^th^ and i^th^+4 residue on ICL2 combined with an additional RMSD (root-mean-square deviation) cutoff. Here, ICL2 was defined as FFAR1 residues 110 to 118. Hydrogen bond detection was performed with standard geometric criterion using the getcontacts software tool (https://getcontacts.github.io/). The RMSD of the ICL2 segment was calculated on ICL2 backbone atoms after aligning this selection to the starting frame, where ICL2 is helical. If 3 or more backbone h-bonds (i,i+4 only) were present in the frame or the RMSD of ICL2 backbone atoms was <2 angstroms, the ICL2 conformation of that frame was classified as helical. In the initial helix, only 4 backbone h-bonds are present and we found the cutoff of 3 helical hbonds was sufficient to accurately capture the state while allowing some flexibility.

We then created two metrics to describe the different ICL2 helical conformations. First, we calculated the angle (about the helical axis) of ICL2 relative to the rest of the receptor. The receptor was aligned on TM1,2,3, and 4. We found that this placed the helical axis of ICL2 approximately perpendicular to the z,y plane. Then, a vector was drawn between atoms 114:OH and 112:CG in the z,y plane. The beginning of this vector was placed at a hypothetical origin, and then the ICL2 angle was calculated as on a typical unit circle. For displayed traces of this angle, we then set the initial angle in the crystal structure as 0 degrees, to provide a convenient point of reference. As a second metric, we calculated the distance between Y114 and TM2 (using the Y114:OH atom to T39:CB atom). This allowed us to detect whether Y114 had moved into or away from the helical bundle.

To calculate fraction of time spent in the PR or NR state, we developed a set of simple thresholds to assign the state. These thresholds were based on the ICL2 angle (*a*) and the Y114 to TM2 distance (*d*) in order to create well separated clusters of states. The PR state was assigned if ICL2 was helical, 45<a<120, and 195-20*a < d < 13. The rotated state was assigned if ICL2 was helical but not in the crystal state. The overall results were robust to small changes in these thresholds.

For the more generalized ICL2 analysis shown in Supp. Fig. 12, we used machine learning library sklearn to perform principal component analysis of the phi, psi backbone dihedrals of ICL2 residues 3.55 to 4.39. We only used frames where ICL2 was helical by applying the RMSD cutoff described previously. For each receptor we calculated new principal components using merged data across available conditions and simulation trials for that receptor.

### Additional simulation analysis

To quantify the vertical shift of TM5 relative to TM4, shown in Fig. 1a and Supp. Fig. 2, we projected the position of a Cα atom on TM5 (residue 190), onto the line connecting the Cα atoms of TM4 residues 130 and 141. We then report the position of the projected point on that line, using the convention that the AP8-free crystal structure is at 0 and positive values indicate a upward shift of TM5 toward the extracellular side of the receptor.

To quantify the frequency of hydrogen bonds, shown in Fig. 2b, we used the getcontacts software tool. Hydrogen bond detection was performed with standard geometric criterion. The frequency of hydrogen bonding interactions for an atom pair was calculated by take the number of simulation frames where a hydrogen bond or water-mediated hydrogen bond between specified atoms was detected, and dividing by the total number of simulation frames. The average frequency of each interaction over the ten simulations for each condition was calculated.

To investigate the effect of AP8 on G-protein conformation, we calculated the displacement of Gα helix 5 relative to the receptor for each condition (shown in Fig. 3c). After aligning receptor TM domains 1-4 to the starting frame, we calculated the RMSD of Gqα residues 519 to 530 for each frame. To measure G-protein internal conformation, we analyzed β6-α5 loop (Supp. Fig. 9). We calculated the end-to-end distance of the loop using residue 504 Cα to residue 508 Cα. The average distance over the five simulations for each condition was calculated. We also calculated the average standard deviation of this quantity.

### IP accumulation assay

Human GPR40 WT and mutants were transiently expressed in HEK293 cells (purchased from American Type Culture Collection (ATCC) and mycoplasma tested). HEK293 cells were grown in Dulbecco’s Modified Eagle Medium (DMEM) containing 10% Fetal Bovine Serum (FBS), 1% penicillin and streptomycin (Life Technologies). 40,000 cells per 100 μl per well were seeded in a 96 well poly-d lysine coated plate. Transfection complexes were prepared by adding 5 ug of plasmid DNA (pcDNA 3.4 TOPO) to 300 μl of optiMEM (Life Technologies), and 18 μl of Fugene HD (Promega) to 300 μl of optiMEM. These two solutions were mixed, incubated at room temperature for 20 min and then 10 μl of this solution was added per well to cells in the 96 well cell culture plate (84 ng plasmid DNA per well). 24 hours post transfection, media was changed to optiMEM. 48 hours post transfection, IP1 accumulation assay was performed. On the day of the experiment, the growth media was removed from the assay plate and 40 μl of IP1 stimulation buffer (Cis Bio IP-one Tb HTRF kit) supplemented with 50 mM LiCl added to each well. Test compounds dissolved in DMSO were serially diluted in half-log increments, starting from 1 mM as 100X, diluted to 10X in stimulation buffer before adding 5 μl per well (final starting concentration 10 µM). Plates were then incubated for 60 min at 37°C. 50 μl of Lysis buffe per well was added to each plate and incubated for 60 min at room temperature. 10 μl of detection buffer (prepared as described in the Tb HTRF kit) is added to each well. The plates are then incubated an additional 1 hr and 30 min at room temperature. After the final incubation, the plates were read in a Perkin Elmer Envision with a method designed for HTRF assays (320 nm excitation, dual emission 615 and 655 nm). For each assay, a standard curve plate in which IP1 is titrated is also included. All fluorescent readings (using the 655/615 nm ratio) are back calculated to a concentration of IP1 using the IP1 standard curve. The percent activity at each concentration of test compound is normalized using the basal activity of WT GPR40 determined in GPR40 WT wells that contained DMSO. The % activation is then plotted versus the concentration of test compound and the dose–response curve fitted to a standard 4-parameter nonlinear regression model using a GraphPad Prism 7. Maximal % activation and EC50 are then determined for each test compound.

### Plasmid construction

WT GPR40 was cloned in a pcDNA3.4 TOPO TA vector with a N-terminal Flag tag. GPR40 mutants were generated using site directed mutagenesis at GENEWIZ.

### Whole cell Ligand Binding Assay

Human GPR40 WT and mutants were transiently expressed in HEK293 cells (purchased from American Type Culture Collection (ATCC) and mycoplasma tested). HEK293 cells were grown in DMEM containing 10% FBS (FBS), 1% penicillin and streptomycin (Life Technologies). 10 million cells were seeded in a 10cm cell culture dish and were transiently transfected with 17 μg of plasmid cDNA (pcDNA 3.4 TOPO) and 53 μl Fugene HD (Promega) in 10 ml of HEK293 media. Following 48 hr incubation at 37°C and 5% CO2, the transiently transfected HEK293 cells were harvested using dissociation buffer TrypLE (Thermo Fisher Scientific) and binding assay buffer. The cells were pelleted (1,000 r.p.m. for 5 min) and resuspended in binding assay buffer (50 mM Tris HCl, 5 mM MgCl2, 1 mM CaCl2, 100 mM NaCl and 0.1% fatty acid free BSA, Sigma, pH 7.4). For the assay, [3H]-labeled P4, AP8 or AP3 (two-fold serially diluted in binding buffer with a top working concentration of 500 nM), and 60,000 cells were added to a 2-ml 96-well master block plate (Greiner Bio-One) in a total volume of 125μl. The plate was incubated for 4 h at room temperature and the assay then harvested onto a GF/C filter plate (PE) that had been presoaked in 0.3% polyethyleneimine (Sigma) using a Packard Filter Mate Harvester. The plate was washed 4× with 1 ml cold wash buffer (50 mM Tris HCl, 5mM MgCl2, 500 mM NaCl, 2 mM EDTA and 0.05% Tween 20), dried for 1 h at 40 °C and then 50 μl of MicroScint 20 (PE) was added to each well. Plates were then read on a TopCount scintillation counter. Nonspecific binding was determined by the addition of 50 times of cold P4, AP8 or AP4 to control wells and was subtracted from all wells of total binding, in which DMSO was added instead of cold compounds. Kd values were determined by a standard one-site specific binding model.

### Expression analyses by Flow cytometry

HEK293 cells transfected with WT or mutant GPR40 plasmids for IP1 assay and or Binding Assay were harvested with TrypLE (Thermo Fisher Scientific), resuspended in optiMEM (Life Technologies) with 5% heat inactivated Fetal bovine serum (HI-FBS) (Gibco). 1 million cells per 100 μl per sample were stained with anti-Flag antibody (Abcam) (1:100) at Room temperature for 2 hours. Cell were washed 3 times with optiMEM+5% HI-FBS, then stained with a secondary antibody Anti Rabbit-Alexa 488 (Cell Signaling) (1:200) for 1 hr at room temperature. Cells were then washed again, resuspended in optiMEM+5%HI-FBS, and read on a Accuri C6 flow cytometer. Median Fluorescence intensities of Alexa-488 of 20,000 cell per sample were collected and used to calculate % of WT where WT intensities were normalized to 100%.

