## Supplementary Information for "A non-canonical mechanism of GPCR activation"

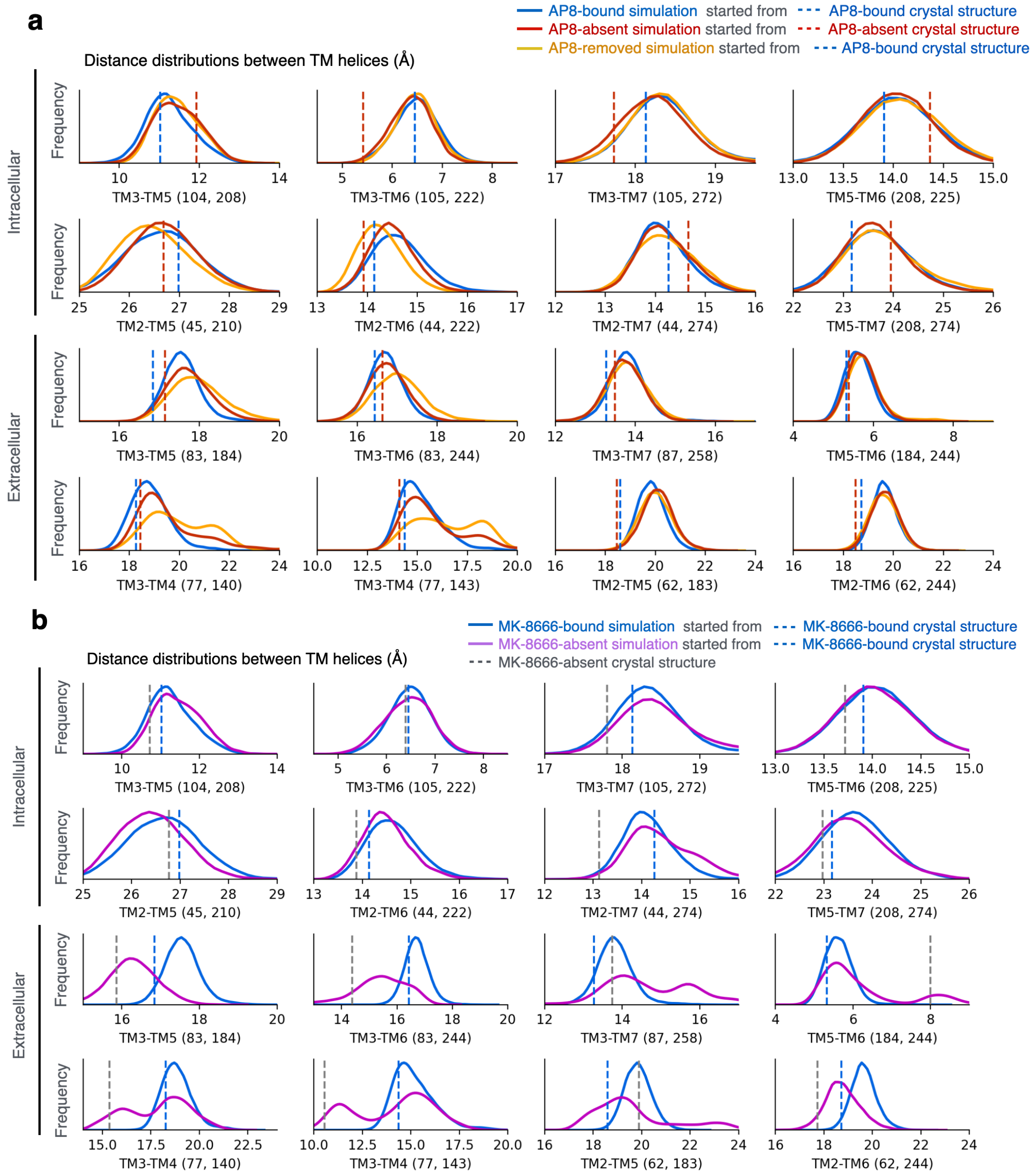


**Supplementary Figure 1. AP8 has minimal effect on the conformation of key transmembrane helices in simulation. (a)** Distributions of distances describing conformation of transmembrane helices under different simulation conditions: blue started from AP8-bound crystal structure (PDB 5TZY), red started from AP8-absent crystal structure (PDB 5TZR), and orange started from AP8-bound crystal structure with AP8 removed. Distances between transmembrane helices were measured using indicated pairs of Cα on the intracellular or extracellular end of each helix. Dotted vertical lines show distances in the initial crystal structures. Plots include simulation frames after 500 ns from 10 independent simulations for each condition. **(b)** As a positive control, we applied the same analysis to simulations with and without FFAR1 partial agonist MK-8666, which binds closer to "canonical" orthosteric site within the helical bundle. MK-8666 does have a substantial effect on key transmembrane helices which shifted by 2-6 angstroms.

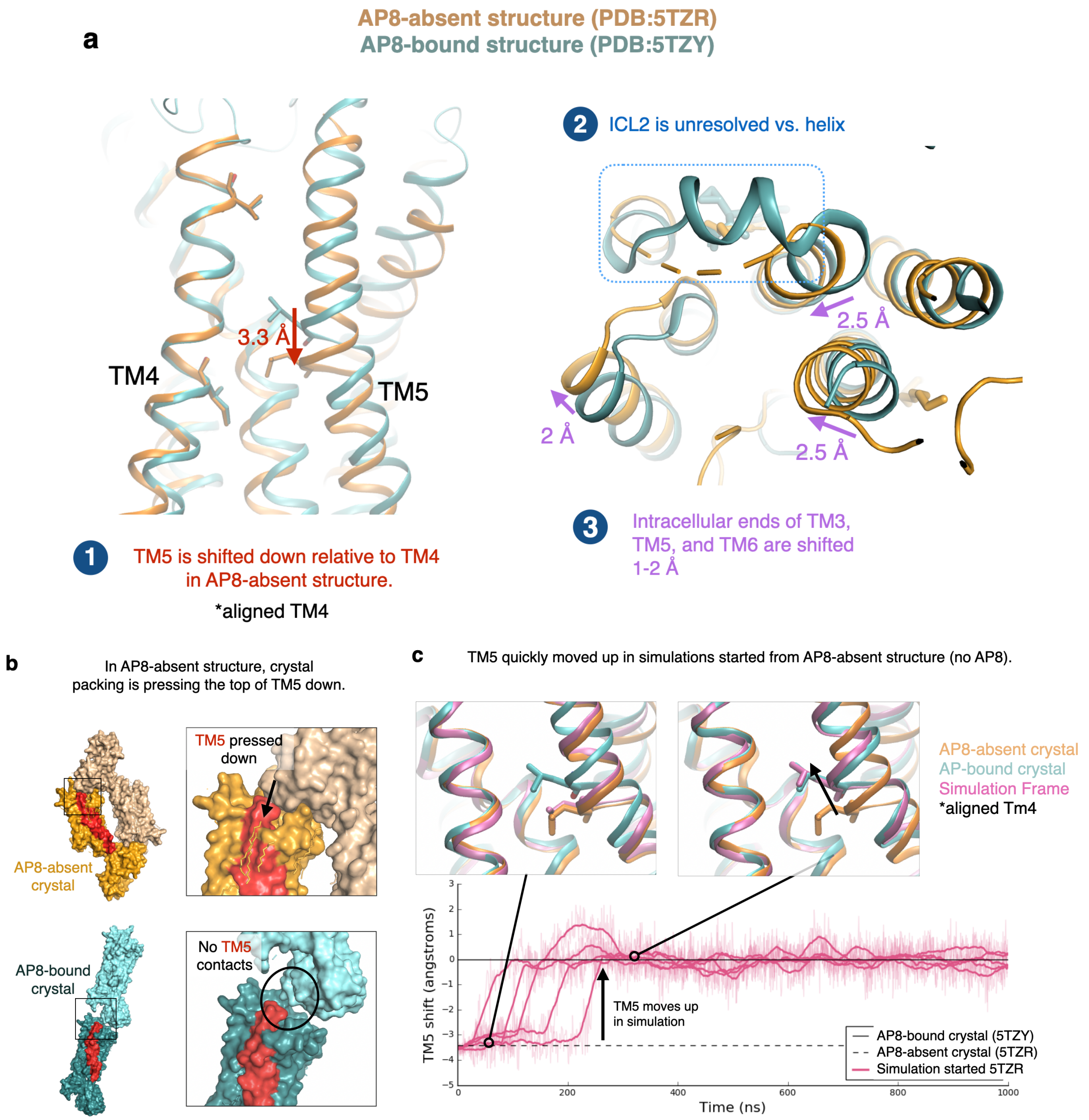


**Supplementary Figure 2. Differences in conformation of transmembrane helices in crystal structures may be a result of crystal packing, not ligand binding. (a)** The conformation of TM5 differs in the AP8-bound and AP8-absent crystal structures, as pictured at left. The receptor is aligned to TM4 to show that TM5 is shifted 3.3. angstroms downward in the AP8-absent crystal structure. The intracellular ends of TM3, 5, and 6 are shifted up to 2.5 angstroms between the AP8-bound and AP8-absent crystal structures, as pictured at right. The receptor was aligned to TM1, 2  3, and 4. **(b)** The AP8-absent crystal structure has a different type of crystal packing, with neighboring subunits pressing TM5 (red) downward. The AP8-bound crystal structure lacks this contact. **(c)** In simulations started from the AP8-absent crystal structure (no AP8 present), TM5 quickly shifts upward to the same conformation as in the AP8-bound crystal structure. This suggests this shift is primarily due to the difference in crystal packing, rather than AP8 binding. Simulation trajectories show the TM5 shift for 5 independent simulations (see methods for details of the metric). Dashed line indicates the distance in the AP8-absent structure while the solid line is the distance in the AP8-bound crystal structure.

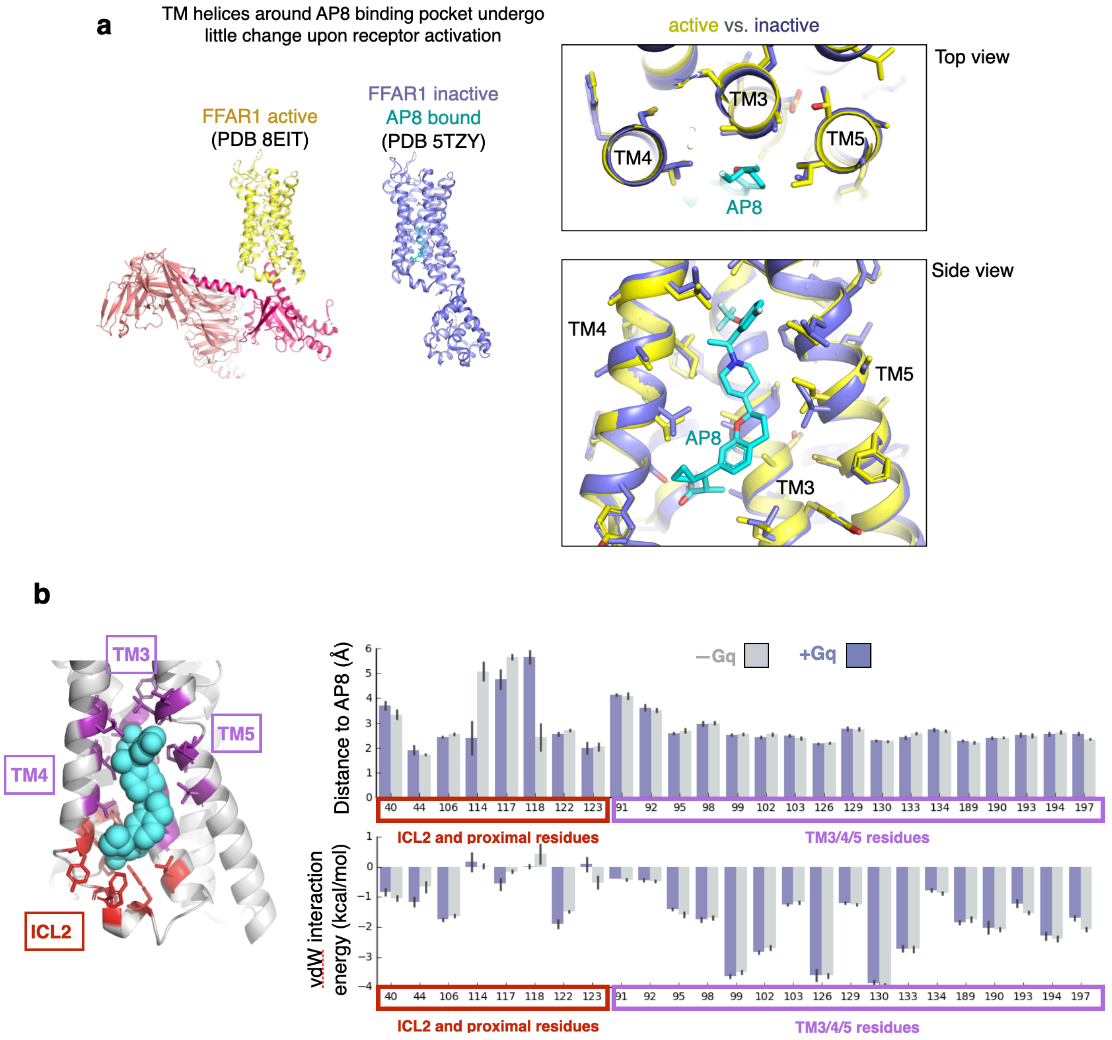


**Supplementary Figure 3. No favorable changes in interactions between AP8 and TM residues, with and without G-protein. (a)** Comparison of cryo-EM structure of the FFAR1-Gq complex (PDB 8EIT, yellow) with the inactive, AP8-bound FFAR1 crystal structure (PDB 5TZY, purple). TM helices surrounding the AP8 binding site undergo little change upon receptor activation; the root mean square deviation of aligned residues shown in images is 0.65 Å (using residues 122-133, 94-106, and 190-201). **(b)** We used in-place docking (Schrodinger's Glide software) to calculate interaction metrics between AP8 and each residue in the binding site, using simulations frames. Two simulation conditions were used: the inactive WT receptor with AP8 bound (light grey) and the active, G-protein complex with AP8 bound (blue) based on our model. Note that our active-state model is very similar to the recently published cryo-EM structure around the AP8 binding site. We used 5 independent simulations per condition with 10 frames (every 100 ns) extracted from each simulation for analysis. The mean values for each metric and each residue were computed for the two conditions. The error bars show the 68% CI.

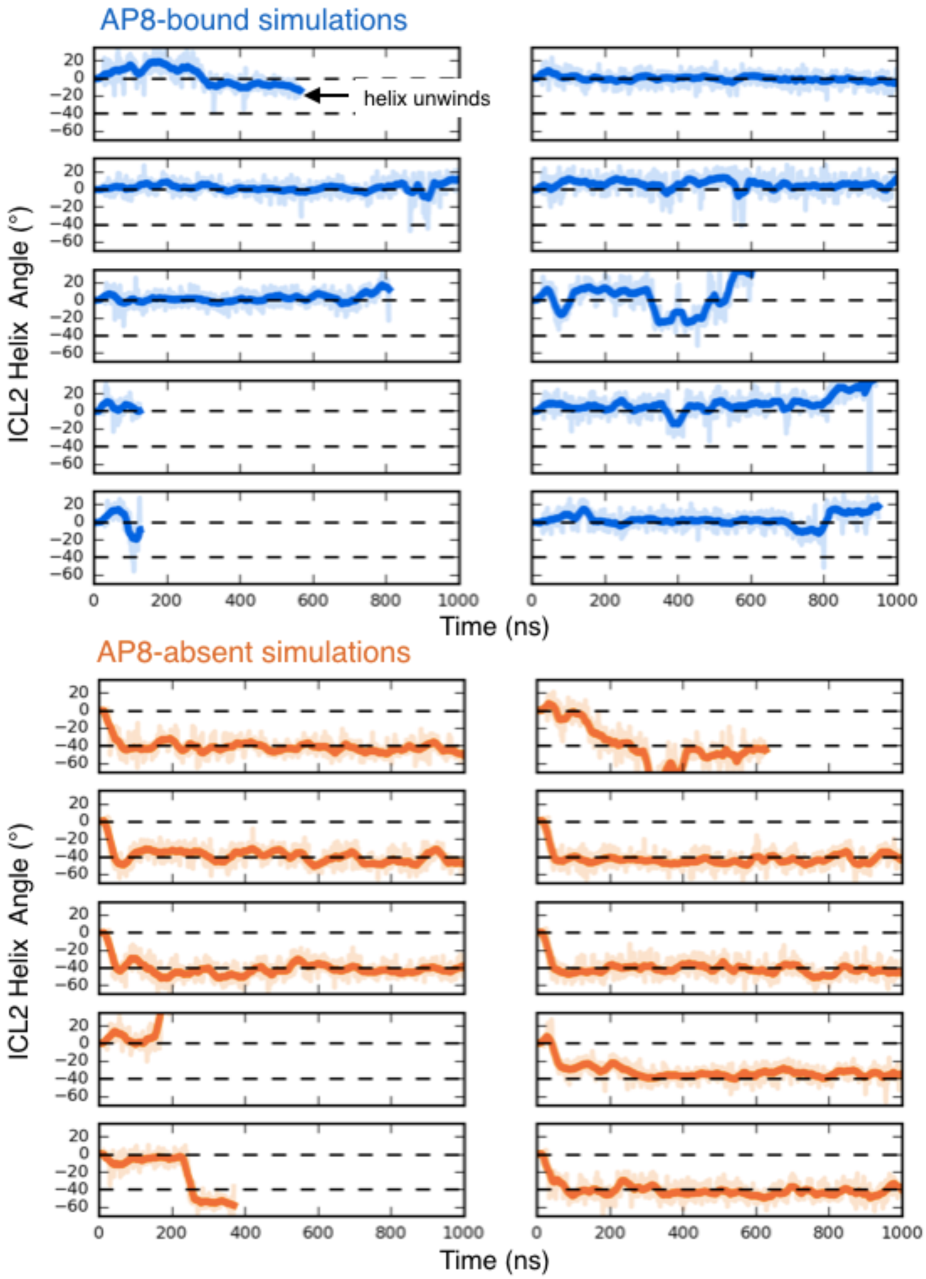


**Supplementary Figure 4: AP8 modulates ICL2 angle in simulations.** The ICL2 helix angle vs. time is shown for each independent simulation trial. The ICL2 angle was measured using the rotation of L112 and Y114 around the ICL2 helix axis for frames where ICL2 remained helical (see methods). All simulations were the same length, though helix unfolding could occur before that (trace ends early). For clarity only the first 1000 ns of the simulations are shown; the average helix unfolding time is 1050 ns with AP8-bound and 3500 ns with AP8-absent. Angles corresponding to the two stable states are labeled with dotted lines. The top line at 0 degrees is the PR state, the bottom line at -40 degrees is the NR state. Simulations of the receptor with AP8 bound (blue) favor the PR state while simulations without AP8 (orange) favor the NR state. All simulations were started from the crystal structure with AP8-bound; AP8 was removed in orange traces. Thick traces are a 15 ns moving average.

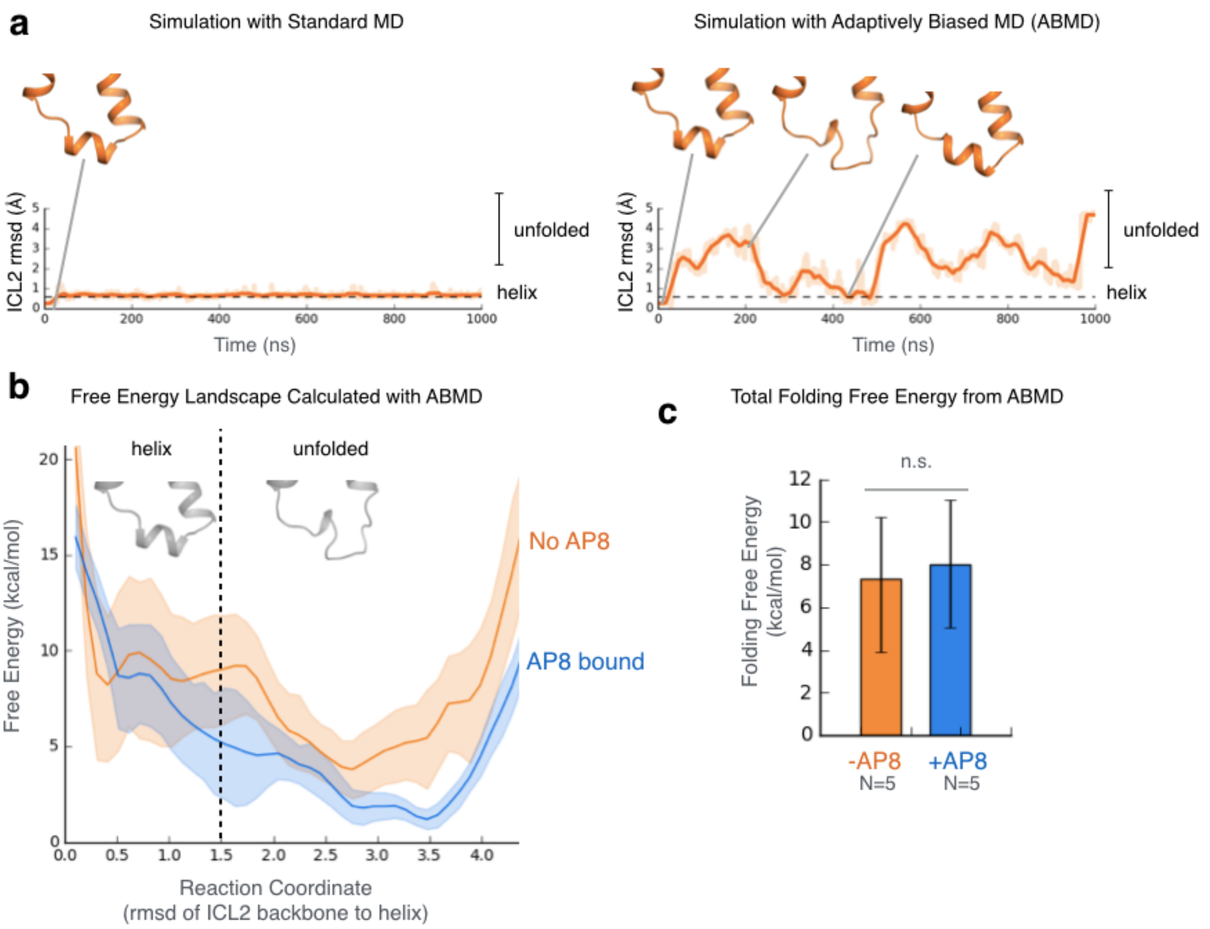


**Supplementary Figure 5. Quantifying folding free energy of ICL2 helix with adaptively biased MD (ABMD).** **(a)** Reaction coordinate chosen was the RMSD (root mean square deviation) of ICL2 backbone to the ICL2 helix. Using ABMD with this reaction coordinate led to more uniform sampling of the reaction coordinate and more frequent transitions between states as expected (right trace). This is in contrast to standard MD (left trace) which shows limited sampling of the conformational space. **(b)** The free energy landscape across the helix folding reaction coordinate was calculated with and without AP8 bound. The results are the average of 5 simulations for each condition. The shaded area shows the standard error of the mean. **(c)** The net folding free energy was not significantly different with and without AP8 bound. To get the net folding free energy difference, we first calculated the relative probability of the folded vs. unfolded states by integrating the free energy landscape from (b) using the partition function, and then transforming the relative probability to an energy difference. Error calculated from min and max curves of the energy landscape.

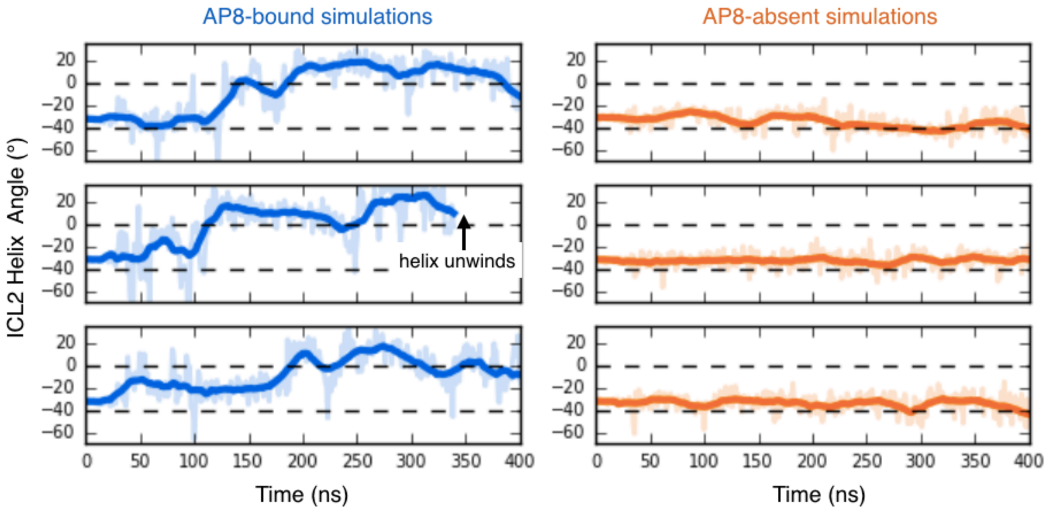


**Supplementary Figure 6: When simulations are started from the rotated ICL2 state, with and without AP8 bound, we observe the same end result.** The ICL2 angle vs. time is shown for each independent simulation trial. Thick traces are a 15 ns moving average. The ICL2 helix angle was measured using the rotation of L112 and Y114 around the ICL2 helix axis (see methods). Angles corresponding to the two stable states are labeled with dotted lines. The top line at 0 degrees is the PR state, the bottom line at -40 degrees is the NR state. Simulations of the receptor with AP8 bound (blue) favor the PR state while simulations without AP8 (orange) favor the NR state. Simulations were started from the NR state, which was modeled into the starting structure using Schrodinger (see methods).

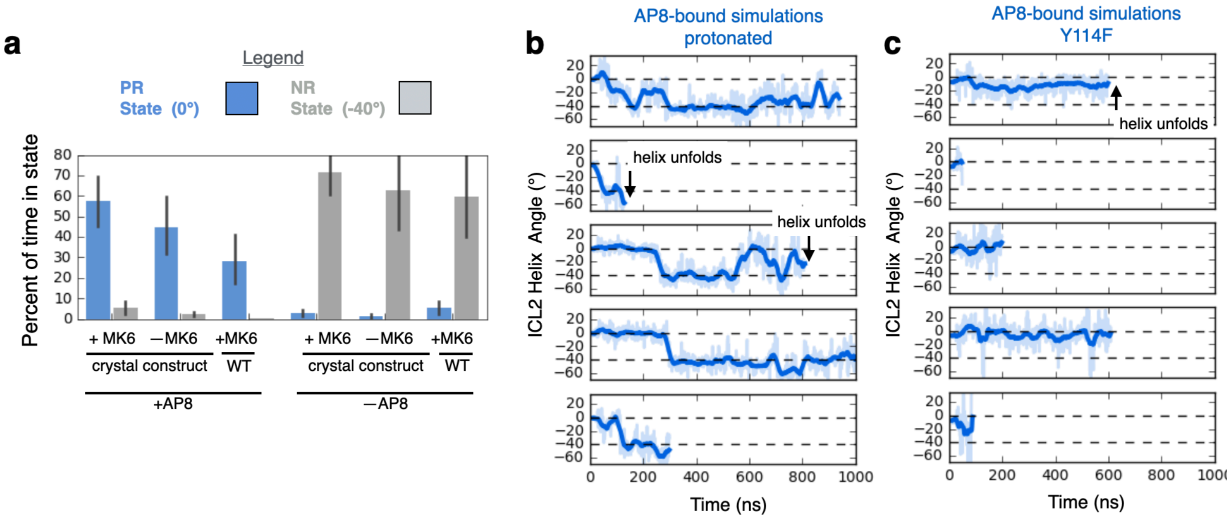


**Supplementary Figure 7. Effect of AP8 on ICL2 across control and perturbed conditions. (a)** Control simulations with partial agonist MK-8666 removed or engineered mutants reversed showed similar results. The major determinant of ICL2 conformation was the presence or absence of AP8. Bars show the mean time spent in each ICL2 helix conformation, over the course of a 1 µs simulation. Error bars show 68% CI and from left to right, N=10, 10, 6, 10, 10, 6. **(b)** Disruption of key hydrogen bond negates AP8's effect on ICL2 conformation. The ICL2 angle vs. time is shown for each independent simulation trial. The top dashed line at 0 degrees is the PR state, the bottom dashed line at -40 degrees is the NR state. AP8's carboxylate group was protonated for the duration of these simulations, disrupting the water-mediated hydrogen bond to ICL2 backbone. This allowed AP8 to adopt the NR state, even with AP8 bound. **(c)** Similarly, Y114 was mutated to phenylalanine, disrupting the hydrogen bond between AP8 and the tyrosine hydroxyl. This hydrogen bond was less important for maintaining the PR state though resulted in faster ICL2 unfolding.

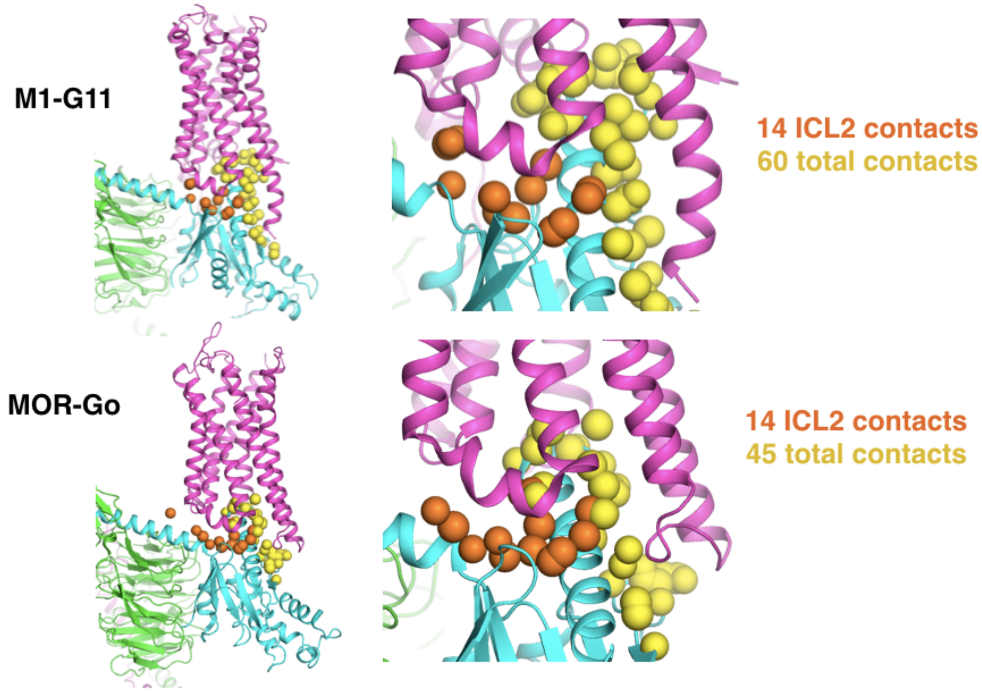


**Supplementary Figure 8. ICL2 forms 25-33% of the interface between the GPCRs and G proteins. (top)** M1 (pink) and G11 (cyan) cryo-EM structure is shown as a cartoon. Spheres show atoms on the receptor that contact the G protein (defined by an atom center-center distance of < 3 angstroms). The contact atoms are colored orange if they are part of ICL2. 23% of contacts occur on ICL2. **(bottom)** MOR (pink) and Go (cyan) cryo-EM structure is shown as a cartoon. 31% of contacts occur on ICL2.

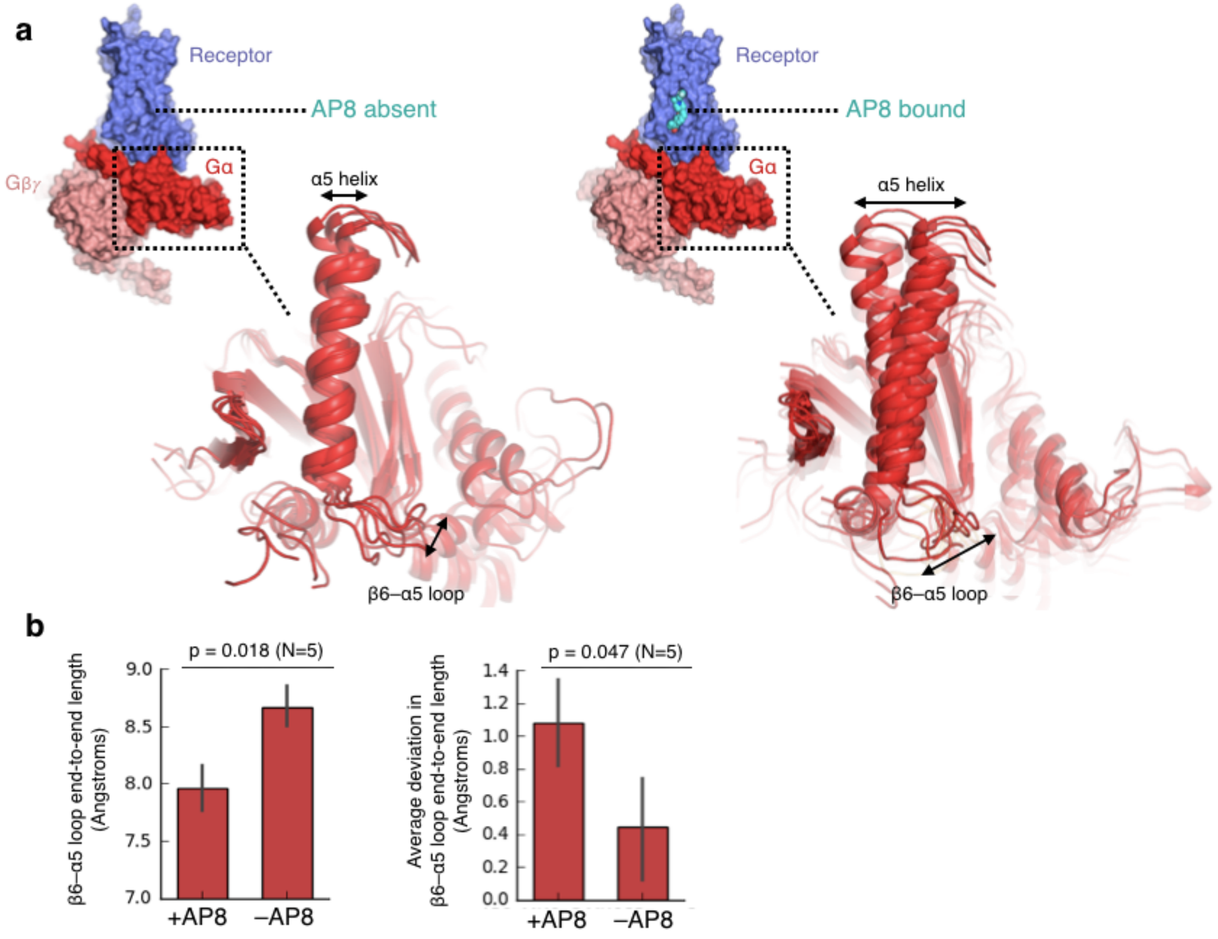


**Supplementary Figure 9. G protein internal conformation and dynamics affected by the presence of AP8. (a)** Cartoon shows an increase in dynamics of α5 helix and connected β6-α5 loop in the presence of AP8. Five frames from different simulations, all taken at the 1000 ns timepoint, are overlaid for each condition. The Gα subunit is aligned to the stable β sheets to better visualize internal motions. **(b)** Comparison of conformation and dynamics with and without AP8 bound, using two metrics. The β6-α5 loop end-to-end length corresponds to length between residue 504 Cα to residue 508 Cα. Left: the end-to-end length of β6-α5 loop is slightly smaller with AP8. Right: the fluctuations in the end-to-end distance increased in the presence of AP8. Data presented as mean from 5 independent simulations for each condition, each 1 μs in length (error bars are 68% CI, P-values calculated using Mann-Whitney U test).

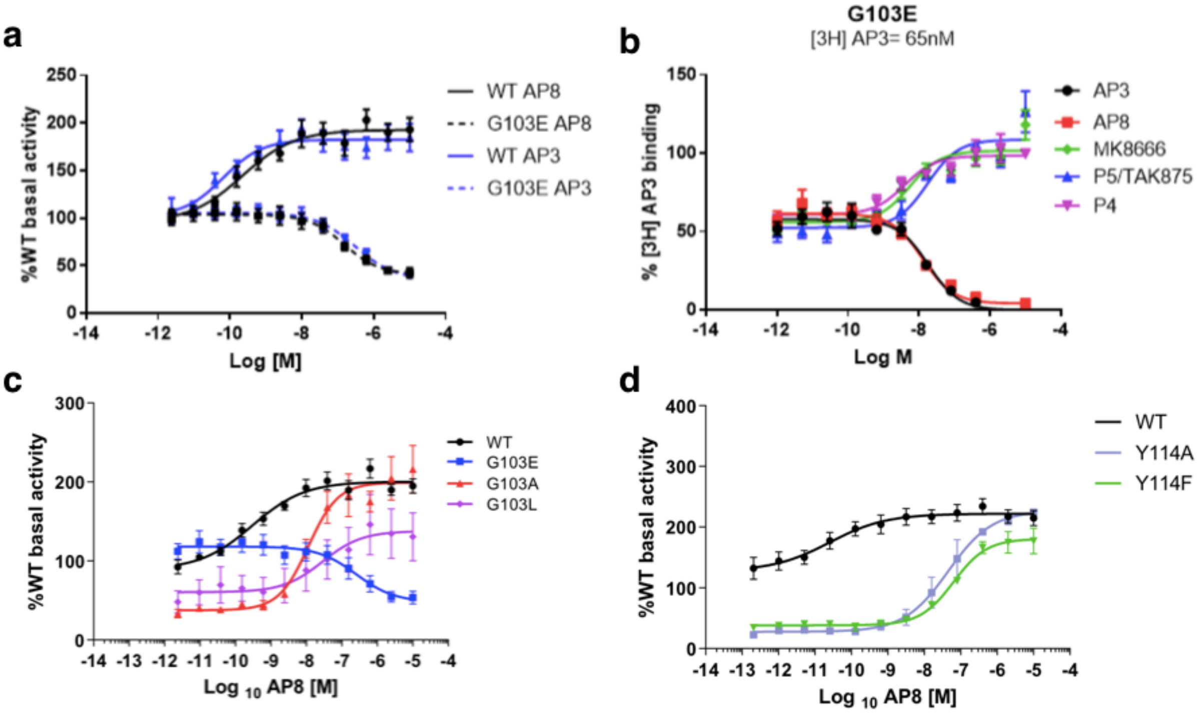


**Supplementary Figure 10. Mutagenesis and binding experiments validate computational results. (a)** G103E converts AP8 to an inverse agonist; consistent behavior was seen in AP3, an analogue of AP8, that binds at the same membrane-facing site. GPR40 activity was monitored in IP1 accumulation assays in HEK293 cells expressing WT or mutant treated with AP8 or AP3. Data is plotted as the % of WT receptor basal activity (cells treated with 1% DMSO), where data points are mean ± S.E.M. from at least N=3 experiments. **(b)** Positive binding cooperativity between AgoPAM AP3 and partial agonists MK-8666, TAK875, and P4, which all bind to the extracellular pocket, is maintained at the G103E receptor. The binding of radiolabeled AP3 was measured at varying concentrations of probe ligand (AP3, AP8, MK-8666, TAK875, and P4). Data points are mean ± S.E.M. from at least N=2 experiments. (**c, d)** Receptor activity was monitored in IP1 accumulation assays in HEK293 cells expressing WT or mutant treated with AP8. Each study was repeated N=2-5 times. **(c)** 103(3.49) mutants produced a range of AP8 efficacies. G3.49E had the lowest *in vitro* efficacy, G3.49L had an intermediate value, and finally G3.49A and WT(G3.49) had the highest values. **(d)** Mutant Y114A and Y114F disrupt the hydrogen bond interaction between Y114 and AP8, contributing to a reduced potency. Calculation of intrinsic efficacy[1,2] indeed showed a 10x decrease for AP8 (P=0.03, t-test), while the effect for MK-8666 was not statistically significant.

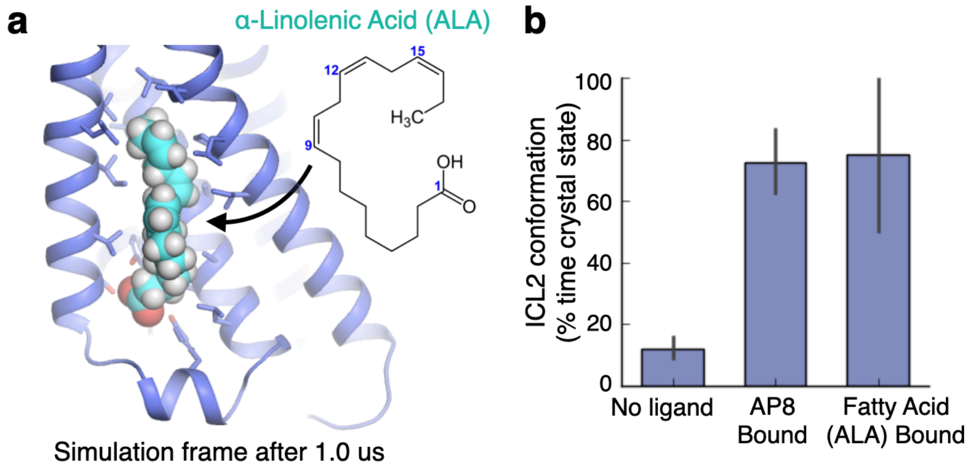


**Supplementary Figure 11. Endogenous fatty acids also bind to and control membrane-facing ICL2 site. (a)** Representative frame from simulation shows that α-linolenic acid (ALA) remains stably bound to the AP8-binding site for at least 1 us. **(b)** Fatty acids exert the same effect on ICL2 conformation as AP8. We measured the fraction of simulation time ICL2 spent in the crystal helix conformation. Error bars show s.e.m. from 5-10 independent simulation trials.

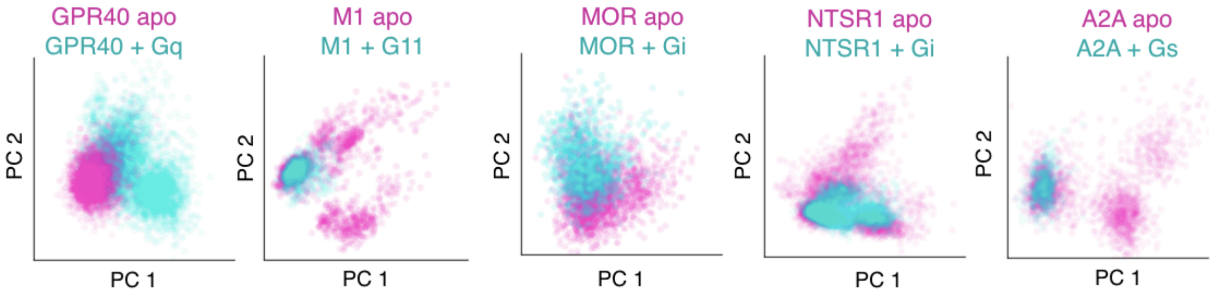


**Supplementary Figure 12. Across diverse receptors, ICL2 helix conformation changes upon formation of receptor-G protein complex.** Principal component analysis of ICL2 psi and phi angles using residues from 3.55 to 4.39. Cyan dots correspond to frames from simulations of receptor-G protein complex and pink dots are from receptor only simulations. Principal components were computed for each receptor independently.

**Supplementary Table 1. Effect of mutations on basal activity and AP8 efficacy.**

| | Membrane Expression (% WT Expression) | Basal Activity (% WT Activity) | AP8 EC50/IC50 (nM) | AP8 Emax (% WT Activity) |
|---|---|---|---|---|
| **WT** | 100 (10) | 105 ± 3 (3) | 0.18 ± 0.14 (6) | 213 ± 17 (6) |
| **G103E** | 53 ± 5 (9) | 103 ± 13 (3) | 150 ± 13 (3) | 42 ± 6 (3) |
| **G103A** | 126 ± 18 (3) | 37 ± 7 (2) | 12 ± 1.3 (2) | 200 ± 43 (2) |
| **G103L** | 91 ± 6 (2) | 63 ± 31 (2) | 33 ± 6 (2) | 136 ± 56 (2) |
| **G103D** | 153 ± 16 (2) | 12 ± 1 (3) | NA | NA |
| **L112A** | 101 ± 9 (4) | 42 ± 10 (2) | 4.6 ± 4 (2) | 333 ± 17 (2) |
| **L112F** | 38 ± 6 (3) | 165 ± 15 (2) | 0.005 ± 0.005 (2) | 575 ± 7 (2) |
| **Y114A** | 86 ± 8 (3) | 31 ± 5 (2) | 70 ± 60 (2) | 222 ± 12 (2) |
| **Y114F** | 102 ± 11 (4) | 38 ± 6 (2) | 130 ± 48 (2) | 184 ± 35 (2) |
| **A116G** | 76 ± 4 (2) | 53 (1) | 0.05 ± 0.04 (2) | 274 ± 10 (2) |

*Data presented as mean ± s.d. (N independent experiments)

**Supplementary Table 2. Effect of mutations on MK-8666 efficacy.**

| | EC50 (nM) | Emax (% WT Activity) |
|---|---|---|
| **WT** | 4.8 ± 1.9 (5) | 202 ± 24 (4) |
| **G103E** | 13 ± 3.6 (5) | 121 ± 10 (4) |
| **G103A** | 22 ± 2.9 (2) | 152 ± 27 (2) |
| **G103D** | 41 ± 16 (3) | 127 ± 35 (3) |
| **G103L** | 5.3 ± 2.2 (3) | 92 ± 31 (3) |
| **Y114A** | 5.0 ± 4.9 (2) | 137 ± 13 (2) |
| **Y114F** | 14 ± 0.1 (2) | 133 ± 20 (2) |
| **A116G** | 2.7 ± 2.3 (3) | 189 ± 1 (3) |

*Data presented as mean ± s.d. (N independent experiments)

**Supplementary Table 3. Binding Affinity of AP8 and analogs at WT and G103E construct**

| | **Construct** | | |
|---|---|---|---|
| **Radioligand** | **GPR40 WT** | **G103E mutant** | **Y114F mutant** |
| [$^3$H]AP8 | 2.0 ± 0.5 (3) | 28.5 ± 1.2 (3) | 29.3 ± 0.5 (3) |
| [$^3$H]AP3 | 3.1 ± 1.8 (2) | 55.6 ± 21.6 (2) | NA |
| [$^3$H]P4 | 0.9 ± 0.4 (3) | 2.3 ± 0.3 (3) | 1.6 ± 0.1 (3) |

*Data presented as mean $K_d$ (nanomolar) ± s.d. (N independent experiments)

**Supplementary Table 4. Simulation conditions**

| # | Name | Description | Trials | Length (us) |
|---|---|---|---|---|
| 1 | AP8-bound 5TZY | Receptor structure from PDB:5TZY with bound ligands | 10 | >2 |
| 2 | AP8-absent 5TZY | Receptor structure from PDB:5TZY, AP8 removed | 10 | >2 |

| | | | | |
|---|---|---|---|---|
| 3 | AP8-bound, MK8-absent 5TZY | Receptor structure from 5TZY, MK-8666 removed | 10 | >2 |
| 4 | AP8-absent, MK8-absent 5TZY | Receptor structure from PDB:5TZY, AP8 removed, MK-8666 removed | 10 | >2 |
| 5 | AP8p-bound | Receptor structure from PDB:5TZY, AP8 protonated | 5 | 1 |
| 6 | AP8-bound, start rotated | Same setup as 1, but ICL2 modeled to start at rotated state (modeled from condition 2 simulation) | 5 | 0.5 |
| 7 | AP8-absent, start rotated | Same setup as 2, but ICL2 modeled to start at rotated state (modeled from condition 2 simulation) | 5 | 0.5 |
| 8 | AP8-bound ABMD | Same setup as 1, but using adaptively biased MD with reaction coordinate as ICL2 helicity | 5 | 1 |
| 9 | AP8-absent ABMD | Same setup as 2, but using adaptively biased MD with reaction coordinate as ICL2 helicity | 5 | 1 |
| 10 | AP8-bound G103E | Same setup as 1, G103E mutant modeled | 10 | >2 |
| 11 | AP8-absent G103E | Same setup as 2, G103E mutant modeled | 10 | >2 |
| 12 | AP8-bound, MK8-absent G103E | Same setup as 3, G103E mutant modeled | 10 | >2 |
| 13 | AP8-absent, MK8-absent G103E | Same setup as 4, G103E mutant modeled | 10 | >2 |
| 14 | AP8-bound, Y114F | Same setup as 1, Y114F mutant modeled | 5 | 1 |
| 15 | AP8-bound, G103 | Same setup as 1, G103 to replace A103 in crystal construct | 5 | 1 |
| 16 | AP8-bound, G103L | Same setup as 1, G103L mutant modeled | 5 | 1 |
| 17 | AP8-absent 5TZR | Receptor structure from 5TZR (AP8-absent crystal structure) | 5 | 1 |
| 18 | AP8-bound, Gq-bound | Active state model of FFAR1 with bound-AP8 and heterotrimeric Gq | 5 | 1 |
| 19 | AP8-absent, Gq-bound | Active state model of FFAR1 with bound-AP8 and heterotrimeric Gq | 5 | 1 |
| 20 | AP8-bound, Gq-bound, G103E | Same as 18, G103E mutant modeled | 5 | 1 |
| 21 | AP8-absent, Gq-bound, G103E | Same as 19, G103E mutant modeled | 5 | 1 |
| 22 | MOR receptor, Gi-bound | Receptor and G-protein complex from PDB: 6DDF | 5 | 1 |
| 23 | MOR receptor | Same structure as 22, but G protein removed | 5 | 1 |
| 24 | M1 receptor, G11-bound | Receptor and G-protein complex from PDB: 6OIJ | 5 | 1 |
| 25 | M1 receptor | Same structure as 24, but G protein removed | 5 | 1 |
| 26 | NTSR1 receptor, Gi-bound | Receptor and G-protein complex from PDB: 6OS9 | 5 | 1 |
| 27 | NTSR1 receptor | Same structure as 26, but G protein removed | 5 | 1 |
| 28 | A2A receptor, Gs-bound | Receptor and G-protein complex from PDB: 6GDG | 5 | 1 |
| 29 | A2A receptor | Same structure as 28, but G protein removed | 5 | 1 |
